# A mycobacterial glycine-rich protein governs actin-based motility

**DOI:** 10.1101/2021.10.01.462822

**Authors:** Norbert S. Hill, Matthew D. Welch

## Abstract

*Mycobacterium marinum*, a close relative of the significant human pathogen *Mycobacterium tuberculosis,* polymerizes host actin at the bacterial surface to drive intracellular movement and cell-to-cell spread during infection. Here, we report the identification and characterization of MirA, the *M. marinum* actin-based motility factor. MirA is a member of the glycine-rich PE_PGRS family of ESX-5-secreted proteins. MirA uses an amphipathic helix to anchor into the mycobacterial outer membrane and, surprisingly, also the surface of host lipid droplet organelles. The glycine-rich PGRS domain in MirA directly binds and activates host N-WASP to stimulate actin polymerization through the Arp2/3 complex, directing both bacterial and lipid droplet actin-based motility. MirA is dissimilar to known N-WASP activating ligands and may represent a new class of microbial and host actin regulator. Additionally, the MirA-N-WASP interaction represents a model to understand how the enigmatic PE_PGRS proteins contribute to mycobacterial pathogenesis.

---

Elucidating how microbial pathogens manipulate host cell pathways and structures during infection has illuminated many aspects of host cell biology. A frequent target of infectious microbes is the actin cytoskeleton, and microbes have developed an array of mechanisms to commandeer actin polymerization for entry, establishing a replicative niche, and egress (*1–4*). A diverse set of pathogens have evolved the ability to polymerize actin directly against the microbial surface to power actin-based motility. This propels the microbe within and between host cells, driving tissue dissemination while also allowing continued access to cytosolic nutrients and avoidance of host immune responses (*5*). *Mycobacterium marinum*, an aquatic pathogen that occasionally infects humans, and a model organism to study the significant human pathogen *Mycobacterium tubercul*osis (*6, 7*), is one such pathogen that undergoes actin-based motility (*8–10*). To polymerize actin, *M. marinum* recruits the host cell nucleation promoting factors WASP and/or N-WASP, which promote actin filament nucleation by binding to and activating the host Arp2/3 complex. However, the bacterial factor governing actin-based motility in *M. marinum* has remained elusive, raising the possibility that *M. marinum* has evolved a distinctive WASP/N-WASP activation strategy.

To identify the *M. marinum* actin-based motility factor and other genes involved in cell-to-cell spread, we assessed >35,000 individual *M. marinum* transposon insertion mutants by fluorescence microscopy for defects in cell-to-cell spread through a confluent monolayer of U2OS host cells. Within the subset of genes that were crucial for cell-to-cell spread without causing a marked growth defect, we focused on *MMAR_3581* (*MMAR_RS17840*) (**Figure 1A**) because its product is known to be secreted across the bacterial envelope (*11*). *M. marinum* is the only mycobacterial species to encode *MMAR_3581* except for a truncated form encoded by the closely related pathogen *Mycobacterium ulcerans*. The genomic region adjacent to *MMAR_3581* exhibits multiple features of horizontal gene acquisition (**Figure S1**) suggesting that *M. marinum* may have acquired *MMAR_3581* from another mycobacterial species. MMAR_3581 is a member of the PE_PGRS protein family, which are translocated through the ESX-5 type VII secretion system (T7SS) (*12*). Consistent with an ESX-5 substrate being involved in *M. marinum* actin-based motility, two separate transposon insertions into *eccA5*, a cytosolic component of ESX-5 were also identified in our screen as having significantly reduced spread (**Figure S2A**).

**Figure 1.**
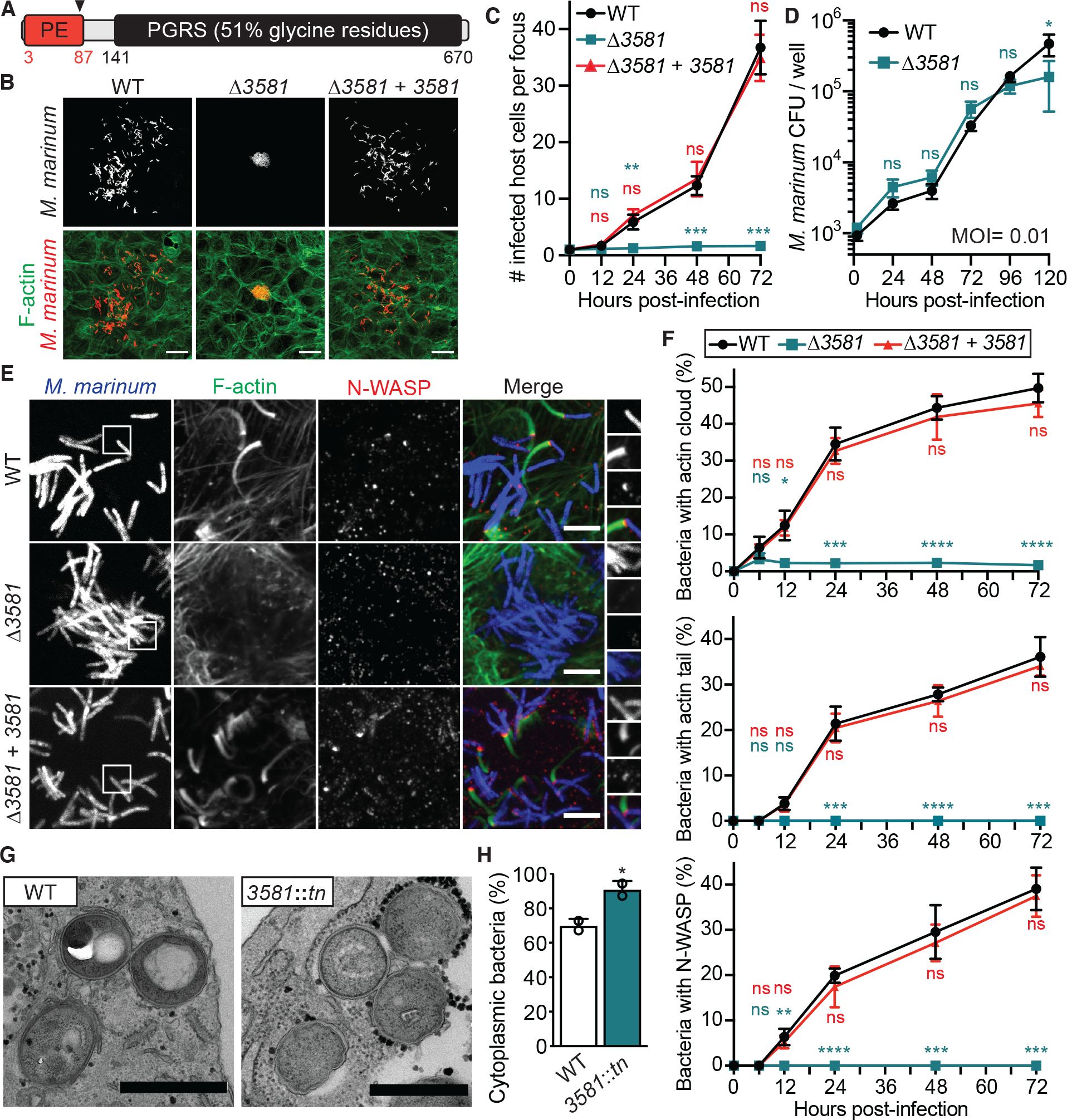
*MMAR_3581* is critical for cell-to-cell spread and actin-based motility in *M. marinum.* (**A**) Schematic of MMAR_3581 (683 amino acids) domain structure. Arrow indicates putative proteolytic processing upon secretion. (**B**) Representative micrographs depicting cell-to-cell spread in a single infectious focus with *M. marinum* strains (red; tdTomato) in U20S host cells (green; F-actin) at 48 hpi. Scale bar is 10 µm. (**C**) Time course graph indicating the number of host U20S cells per infectious focus during infection of indicated *M. marinum* strains. (**D**) Growth curve of *M. marinum* strains during infection of U20S host cells. (**E**) Representative micrographs depicting ability of indicated *M. marinum* strains (blue; EBFP2) to recruit N-WASP (red; *α*-N-WASP) and stimulate actin polymerization (green; 488 phalloidin) in U20S host cells at 48 hpi. Scale bar is 5 µm. (**F**) Time course graphs of *M. marinum* WT stains colocalized with either actin clouds, actin tails, or N-WASP. (**G**) Representative transmission electron micrographs of infected U20S cells to assess vacuolar or cytoplasmic *M. marinum* localization at 60 hpi. Scale bar is 0.5 µm. (**H**) Quantification of cytoplasmic *M. marinum* strains during infection (n = 2). For (**C**), (**D**), (**F**), and (**H**) data is mean ± SD (unpaired t-test); n = 3 unless otherwise indicated.

We generated a clean *MMAR_3581* deletion strain and assessed for its ability to spread through a monolayer of U2OS cells by measuring the number of cells infected per infectious focus. Wild-type *M. marinum* spread efficiently between host cells over a time course of 72 h, however the *MMAR_3581* deletion strain failed to spread (**Figures 1B and 1C**). The defective spread was not due to diminished replication, as the Δ*MMAR_3581* bacteria had indistinguishable growth compared to wild type in broth and during infection of either U2OS or mouse bone marrow-derived macrophages (BMDMs) (**Figures 1D, S2B, and S2C**). Further, complementing the Δ*MMAR_3581* strain with a chromosomally integrated copy of *MMAR_3581* driven by its native promoter restored cell-to-cell spread (**Figures 1B and 1C**). Thus, MMAR_3581, an ESX-5 translocated protein, is necessary for *M. marinum* cell-to-cell spread.

We next tested if the Δ*MMAR_3581* mutant was defective for actin-based motility or a different step in cell-to-cell spread (e.g., phagosome escape or protrusion formation). U2OS cells were infected with wild-type, Δ*MMAR_3581*, or *MMAR_3581* complemented bacteria, then assessed for association with actin clouds, actin tails, and N-WASP over the course of infection. Wild-type *M. marinum* and the complemented strain recruited N-WASP and generated actin tails by ∼12 h post infection (hpi) with a frequency that steadily increased to 36% of the bacterial population by 72 hpi (**Figures 1E and 1F**). Conversely, Δ*MMAR_3581* bacteria were unable to recruit N-WASP and failed to polymerize actin throughout infection. Further, Δ*MMAR_3581* bacteria also failed to undergo actin-based motility in BMDMs (hematopoietic cells that express both WASP and N-WASP whereas U2OS cells only express N-WASP), inferring that *MMAR_3581* is necessary for recruiting both WASP and N-WASP (**Figures S2D and S2E**). Over-expressing *MMAR_3581* >9- fold prompted an earlier timing of actin-based motility (**Figures S2F and S3A**), however the increased MMAR_3581 levels engendered bacterial cell filamentation that caused reduced actin-based motility and spread at later time points (**Figures S2F-I**). Further, the Δ*MMAR_3581* bacteria were not deficient in actin-based motility due to a lack of cytosolic access, as we observed the *MMAR_3581*::*tn* mutant bacteria in the cytosol as frequently as wild type when assessed by transmission electron microscopy (**Figures 1G and 1H**). Altogether, these data demonstrate *MMAR_3581* is necessary to recruit WASP/N-WASP to stimulate actin-based motility during infection. As such, we have designated this gene as *<u>m</u>ycobacterial intracellular rocketing <u>A</u>* (*mirA*).

Previous work demonstrated that that *M. marinum* only initiates actin-based motility in response to the host cell environment (*8*). To examine whether MirA expression is prompted by infection, *M. marinum* grown in either broth or RAW 264.7 murine macrophage cells were probed for MirA expression using an antibody raised against purified MirA. MirA expression was only observed during infection of host cells and coincided with the timing of actin-based motility **(Figures 2A and S3B)**, consistent with a direct role in this process. We next investigated the localization of MirA during infection of U2OS cells. Our MirA antibody was unable to localize MirA by immunofluorescence microscopy, so instead we chromosomally-integrated a gene expressing MirA tagged with a FLAG epitope (MirA-FLAG) driven by its native promoter (**Figure S3A**). MirA was observed at the bacterial surface starting at 8 hpi, and the frequency of MirA surface localization increased over time (**Figures 2B-D**). Of the MirA-positive bacterial population at 72 h, 63% had actin tails and 71% had N-WASP that colocalized with MirA (**Figure 2E**). We observed that the distribution of MirA on the bacterial surface dynamically transitioned over time from a mostly punctate distribution at early time points (8 hpi) to a single polar distribution at later time points (**Figures 2F and 2G**). Together, these data demonstrate that MirA is expressed and secreted in response to infection, and over time it localizes primarily to a single bacterial pole at which N-WASP is recruited and actin is polymerized.

**Figure 2.**
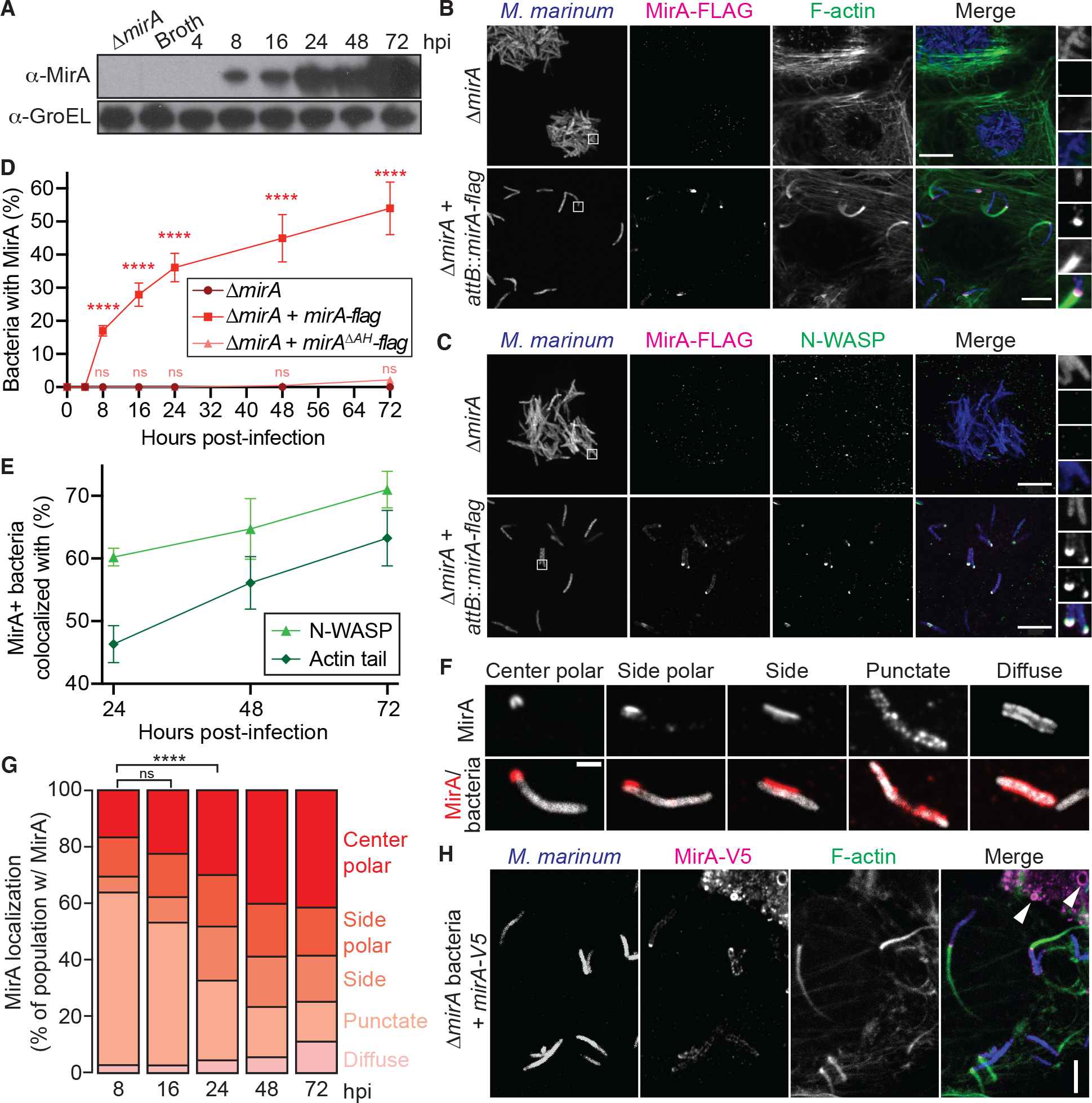
MirA is expressed and secreted during infection, then colocalizes with N- WASP and F-actin. (**A**) Representative immunoblot of MirA expression in broth or during infection of RAW 264.7 cells. GroEL is shown as the loading control. (**B** and **C**) Endogenously-expressed MirA-FLAG (magenta; *α*-flag) with (**B**) F-actin (green; 488 phalloidin) or (**C**) N-WASP (green; *α*-N-WASP) colocalization at the *M. marinum* (blue; EBFP2) surface during infection of U2OS cells. White box indicates zoom panel and scale bar is 5 µm. (**D**) Quantification of indicated *M. marinum* strains with MirA surface localization during infection of U20S cells. Data is mean ± SD; n = 3 (unpaired t-test). (**E**) Percent of MirA-positive bacteria colocalized with either N-WASP or an actin tail over the course of infection. Data is mean ± SD; n = 3. (**F**) Representative micrographs depicting the five categories of MirA-FLAG localization on the bacterial surface during infection of U20S cells. (**G**) Time course of MirA-Flag localization on the bacterial surface during infection of U20S cells. Data is mean; n = 3 (chi-square test). (**H**) Ectopically-expressed MirA-V5 (magenta; *α*-V5) localization with Δ*mirA* bacteria (blue; tdTomato) and F-actin (green; Phalloidin-iFluor 405). Scale bar is 3 µm and white arrows indicate transfected MirA localizing to eukaryotic lipid droplet organelles.

Next, we tested whether ectopically supplied MirA could restore *M. marinum* actin-based motility of Δ*mirA* bacteria. U2OS cells were transfected to express full-length MirA or a variant lacking the PE family domain (MirA^ΔPE^; PE domain, residues 1-87), which is likely cleaved off upon secretion (**Figure S3A**) (*11, 13*). Host cells were subsequently infected with mCherry-expressing Δ*mirA* bacteria and stained for MirA and F-actin at 48 hpi. Ectopically expressed MirA and MirA^ΔPE^ both localized to the bacterial surface and restored actin polymerization (**Figures 2H and S3C-E**). In some cases, MirA concentrated at a bacterial pole and promoted the assembly of an actin comet tail. This indicates that MirA, without its PE domain, can bind to the bacterial surface and has the ability to complement Δ*mirA* bacteria *in trans*.

Curiously, in non-infected eukaryotic cells, ectopically expressed MirA had a distinctive localization to spherical structures (**Figure 2G**, white arrow). We determined the MirA- coated spheres to be lipid droplets (**Figure 3A**), eukaryotic organelles central to lipid and energy homeostasis and hubs for signaling and host defense (*14, 15*). Strikingly, in cells expressing MirA we observed robust actin polymerization at the lipid droplet surface and the formation of actin comet tails (**Figure 3A**). To explore whether MirA was inducing actin-based movement of lipid droplets, MirA was expressed in a U2OS cell line stably expressing F-tractin-mCherry (a fluorescent marker of F-actin) and observed by live-cell imaging. Remarkably, MirA-coated lipid droplets exhibited sustained actin-based motility (**Figure 3B; Movies S1 and S2**). The movement velocity (µm/min), efficiency (final displacement/total distance traveled), and actin tail lengths (µm) were compared between *M. marinum* and lipid droplets. Lipid droplets moved a third slower (mean 13 versus 18 µm/min), more efficiently (mean 0.7 versus 0.5), and had shorter actin tails (mean 3.0 versus 3.7 µm) relative to *M. marinum* (**Figure 3C**). Thus, movement properties were comparable, but differed for each parameter. To assess whether MirA-coated lipid droplets could spread from cell-to-cell, we used a mixing experiment in which cells expressing a plasma membrane marker (RFP-CAAX) were transfected to express MirA, then co-plated with unmarked, non-transfected cells. Remarkably, we observed 5-10% of MirA-coated lipid droplets were found in protrusions of the plasma membrane or in recipient neighboring cells (**Figures 3D and 3E**). Together, these data indicate that MirA is sufficient to induce actin-based motility and cell-to-cell spread of bacteria and lipid droplet organelles.

**Figure 3.**
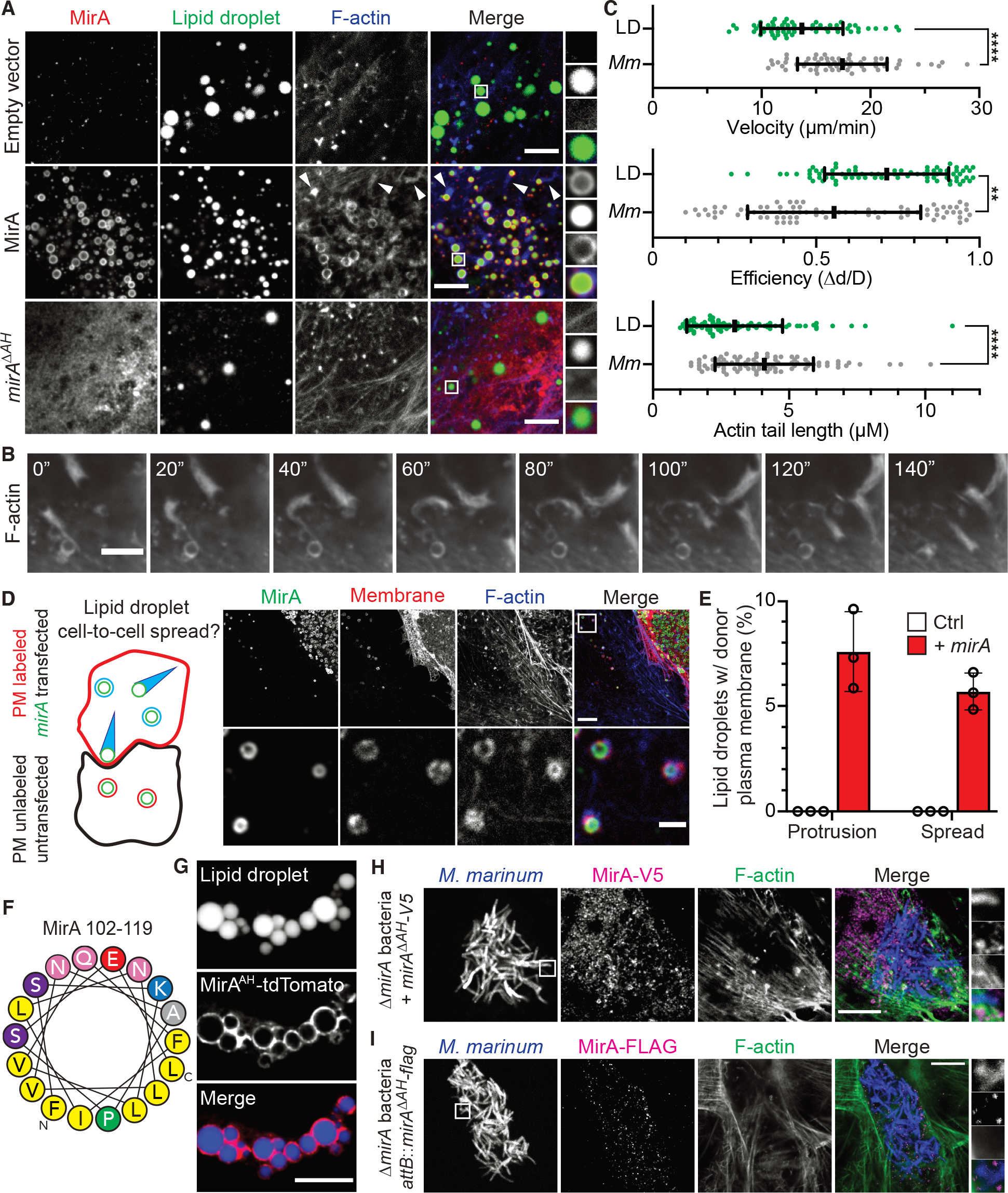
MirA localizes to eukaryotic lipid droplets using an amphipathic helix and stimulates actin rocketing. (**A**) Representative micrographs of ectopically expressed MirA (red; *α*-MirA) in U20S cells localizing to lipid droplet organelles (green; BODIPY 493/503) and induces actin polymerization (blue; Phalloidin-iFluor 405). Scale bar is 3 µm. Arrows indicate actin comet tails. (**B**) Live-imaging snapshots of *mirA^ΔPE^* transfected U20S cells expressing the F-actin marker F-tractin-mCherry. Scale bar is 3 µm. Accompanies **Movie S1**. (**C**) Velocity, efficiency of movement, and actin tail length measurements of *M. marinum* (*Mm*; gray) versus lipid droplet (LD; green) actin rocketing. (**D**) MirA (green; *α*-MirA) coated lipid droplets spread from donor *mirA*-transfected A549 cells expressing a plasma membrane marker (red; mCherry-CAAX) to unmarked and untransfected recipient A549 cells. Bottom panel corresponds to boxed region in the merge panel. Scale bar is 5 µm (upper panel) and 2 µm (lower panel). (**E**) Quantification of MirA-coated lipid droplets from donor cell either in a protrusion or spread vacuole into a recipient cell. (**F**) A putative amphipathic helix in MirA (image generated by HELIQUEST (*16*)). (**G**) The MirA amphipathic helix fused to tdTomato is sufficient to localize to eukaryotic lipid droplet organelles (green; BODIPY 493/503). (**H**) Ectopically expressed MirA^ΔAH^ (magenta; *α*-V5) is unable to localize to the surface of *M. marinum* (blue; expressing tdTomato) and does not promote actin polymerization (green; Phalloidin-iFluor 405). Scale bar is 5µm. (**I**) An endogenously expressed amphipathic helix MirA^ΔAH^ mutant (magenta; *α*-Flag) is unable to localize to the surface of *M. marinum* (blue; EBFP2) to promote actin polymerization (green; 488 phalloidin). Scale bar is 5µm.

To understand how MirA localizes to the surface of lipid droplets and *M. marinum*, we examined MirA for potential membrane targeting sequences. Using HELIQUEST, an online amphipathic helix prediction algorithm (*16*), a putative amphipathic helix was identified in the linker region between the PE family and PGRS domains (AH domain: residues 102-119) (**Figure 3F)**. When the putative amphipathic helix was fused to tdTomato (MirA^AH^-tdTomato) and expressed in U2OS cells, MirA^AH^-tdTomato localized to the surface of lipid droplets (**Figure 3G**). Conversely, deleting the amphipathic helix from MirA (MirA^ΔAH^) compromised lipid droplet targeting and resulted in diffuse cytoplasmic localization without affecting protein levels (**Figures 3A and S3C**). The role of the amphipathic helix was also examined in localization of MirA to the bacterial surface. Unlike wild-type MirA (**Figure 2B**), MirA^ΔAH^ ectopically expressed in U2OS cells was unable to localize to the surface of Δ*mirA* bacteria (**Figures 3H, S3C, and S3D**). Likewise, MirA^ΔAH^ endogenous expressed in bacteria was unable to localize to the bacterial outer membrane and exhibited punctate localization in the host cytoplasm (**Figures 2C**, **3I**, **and S3A**). Bioinformatic analysis of both *M. marinum* and *M. tuberculosis* PE_PGRS proteins, which are largely thought to localize to the bacterial surface (*11, 17–22*), revealed that many score well for encoding a putative amphipathic helix in their linker region (**Figures S4A and S4B**). Further, we find another *M. marinum* PE_PGRS protein MMAR_2645 also localizes to lipid droplets when ectopically expressed in eukaryotic cells (**Figures S4C and S4D**), although it does not promote their actin-based motility. Together, these data demonstrate that MirA uses an amphipathic helix to insert into phospholipid monolayers of both host lipid droplet organelles and the mycobacterial outer membrane, and that amphipathic helices are likely a common feature of mycobacterial PE_PGRS proteins.

To determine if the molecular mechanism of actin-based motility is similar between MirA- coated lipid droplets and *M. marinum*, we stained for N-WASP and the Arp2/3 complex in *mirA* transfected U2OS cells. We observed that both N-WASP and the Arp2/3 complex localize to lipid droplets, though only in cells with MirA expression (**Figure 4A**). We next sought to leverage this system to determine if MirA interacts with N-WASP and explore other interactions with host proteins on lipid droplets using a non-biased affinity purification-mass spectrometry (AP-MS) approach. MirA^ΔPE^ tagged with a Twin-Strep-tag (MirA^ΔPE^-Strep) was expressed in HEK293 cells, then purified by Strep-Tactin affinity chromatography. Differences in the mass spectrometric profile of proteins purified from cells transfected with *mirA-strep* versus an empty plasmid control were assessed by the <u>m</u>ass spectrometry interaction <u>st</u>atistics algorithm (MiST), a computational tool for scoring protein-protein interactions (*23*). MirA-Strep eluates were enriched for N-WASP, three members of the WASP-interacting protein family (WIPF1, WIPF2, and WIPF3), and the canonical N-WASP activator CDC42 (**Figures 4B and S5A**) (*24, 25*). (This approach also identified a possible Prohibitin-Prohibitin2-Erlin1-Erlin2 complex; however, these proteins are not known to be involved in actin cytoskeletal dynamics and were not examined further.) N-WASP, CDC42, and WIPF2 were additionally detected in immunoprecipitates from cells expressing MirA-Strep (**Figure S5B**). The MirA^ΔAH^ mutant was still able to pull down N-WASP, demonstrating that MirA could mediate this interaction in cells without being affixed to the surface of lipid droplets (**Figure S5C**). Further, an inverse approach using N-WASP-Strep as an affinity matrix pulled down MirA when they were co-expressed (**Figure S5D**). Together, these data show that MirA is sufficient to recruit N-WASP and N-WASP-interacting proteins to the lipid droplet surface.

**Figure 4.**
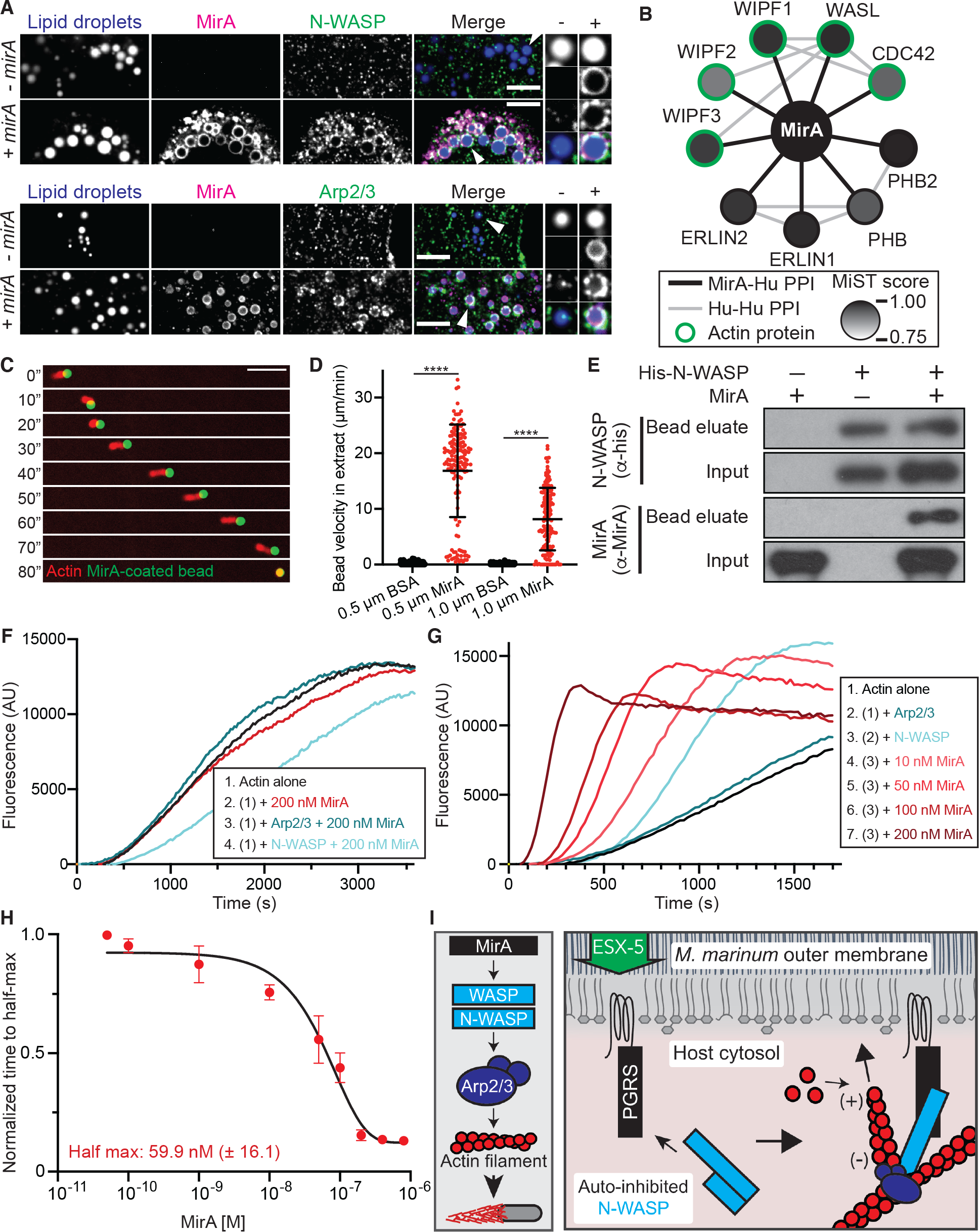
MirA interacts with N-WASP and activates N-WASP to stimulate actin polymerization through the Arp2/3 complex. (**A**) Transfected MirA (red; *α*-MirA) localizes with N-WASP (upper panel) (green; *α*-N-WASP) or Arp2/3 (lower panel) (green; *α*-P34) at the surface of lipid droplet organelles (blue; BODIPY 493/503). Scale bar is 3 µm and the white arrow in merge panel indicates highlighted example. (**B**) A schematic of the MirA interactome determined through an AP-MS approach detecting protein-protein interactions (PPI) determined using the MiST algorithm(*23*). Black lines indicate MirA- human protein interactions, grey lines indicate human-human interactions (via STRING (*49*)), and actin cytoskeletal proteins are circled in green. (**C**) Time-lapse micrographs of a 1 µm fluorescent polystyrene bead coated with native MirA in *Xenopus laevis* egg extract with rhodamine-actin. Scale bar is 5 µm. (**D**) The average velocity (µm/min) of either 0.5 µm or 1 µm beads coated in either BSA or MirA in *Xenopus laevis* egg extract. Middle bar is mean ± SD (unpaired t-test). (**E**) A pulldown assay using purified 6xHis-N-WASP as bait and purified MirA as prey probed with *α*-His or *α*-MirA by immunoblot. (**F**) Representative pyrene actin polymerization reactions with MirA in tandem with actin alone (1 µM, 10% labeled), actin and the Arp2/3 complex (50 nM), or actin and N-WASP (200 nM). (**G**) Representative pyrene actin polymerization reactions with increasing MirA concentrations combined with actin (1 µM, 10% labeled), the Arp2/3 complex (50 nM) and N-WASP (200 nM). (**H**) The time to half-maximum fluorescence of the polymerization curves normalized to N-WASP and the Arp2/3 complex without MirA. Data is mean ± SD; n = 3, and the data set was fit to estimate the concentration at which half-maximum activity is observed. (**I**) A model of *M. marinum* actin-based motility whereby MirA is translocated by ESX-5, inserts into the outer membrane though an amphipathic helix, then recruits and directly activates host N-WASP to stimulate actin polymerization against the bacterial surface for motility.

To initially assess the biochemical activity of MirA in actin-based motility, we purified recombinant MirA protein, bound it to the surface of polystyrene beads, and incubated MirA-coated beads in a *Xenopus laevis* egg extract supplemented with rhodamine-labeled actin. The MirA-coated beads initiated actin-based motility and moved at a similar velocity to *M. marinum* or MirA-coated lipid droplets (0.5 µm beads had an average velocity of 16.9 µm/min) (**Figures 4C and 4D**). To further determine if MirA and N-WASP directly interact, a pulldown assay was employed using recombinant purified MirA and 6xHis-N-WASP. MirA was pulled down only when pre-incubated with N-WASP, suggesting the two proteins bind directly (**Figure 4E**). To test if MirA activates N-WASP, thereby allowing it to bind the Arp2/3 complex to stimulate actin nucleation, we tested their activity in actin polymerization assays. Purified MirA had no effect on the kinetics of actin assembly when added to actin alone, or with either N-WASP or the Arp2/3 complex (**Figures 4F and S5E**). These results suggest that MirA is neither an actin nucleator nor an Arp2/3-binding nucleation promoting factor. However, when added together with N-WASP and the Arp2/3 complex, MirA accelerated actin assembly in concentration-dependent manner (>7.5-fold increase relative to N-WASP alone) (**Figure 4G**) with 60 nM required for half-maximum activity (**Figure 4H**). Thus, MirA promotes actin assembly by directly binding and activating N-WASP to stimulate Arp2/3-dependent actin filament nucleation.

Our work reveals that MirA is the *M. marinum* actin-based motility factor. We demonstrate that in response to infection, MirA is expressed, translocated, and anchored into the bacterial outer membrane though an amphipathic helix. There, MirA directly binds and activates N-WASP to stimulate actin filament nucleation through the Arp2/3 complex, which drives bacterial intracellular motility and cell-cell spread (see a model in **Figure 4I**). MirA is a member of the PE_PGRS protein family, the largest group of proteins exported by virulent mycobacteria. Yet, the contribution of individual PE_PGRS proteins to infection is poorly understood, in part because these proteins lack sequence homology with other proteins in existing databases and have been recalcitrant to genetic or biochemical manipulation (*26, 27*). PE_PGRS proteins are thought to primarily localize to the mycobacterial outer membrane, though how this localization is achieved has been an open question (*26*). We demonstrate that MirA contains an amphipathic helix between the PE and PGRS domains that is necessary and sufficient for localization to the bacterial surface. We also find that amphipathic helices are a common feature among the PE_PGRS protein family, thereby establishing a mechanism by which PE_PGRS proteins can be targeted to the mycobacterial outer membrane. Further, we hypothesize that the bioinformatic prediction of amphipathic helices (**Figures S4A and S4B**) may enable prediction of PE_PGRS protein localization to the bacterial surface.

Surprisingly, we find that the amphipathic helix can also target MirA to the surface of lipid droplet organelles, also a phospholipid monolayer, when ectopically expressed in eukaryotic cells. There, MirA stimulates actin polymerization that propels lipid droplets by actin-based motility, analogous to bacterial movement. This raises the possibility that MirA could be targeted to lipid droplets during infection. Recent work has shown that a host lipid droplet protein containing an amphipathic helix is transferred to the *M. marinum* surface during infection (*28*), suggesting that targeting of bacterial proteins to host lipid droplets may also occur. Further work will be necessary to determine if MirA and/or other PE_PGRS proteins are transferred to the host lipid droplet surface to impact infection, for example by disrupting host immune signaling (*29, 30*) or enabling nutrient acquisition.

PE_PGRS proteins are characterized by the PGRS domain, which is enriched for glycine residues, specifically GGX repeats. For instance, MirA has a ∼529 amino acid PGRS domain composed of 51% glycine residues with 102 GGX sequences. Yet, it has been unclear how these glycine-rich sequences contribute to infection and whether they target specific host proteins (*31, 32*). A handful of the PE_PGRS proteins have been reported to contribute to functions related to host-pathogen interaction, such as suppressing autophagy (*33, 34*), inhibiting lysosomal fusion (*35*), and antagonizing signaling from Toll-like receptor proteins (*20, 36–38*). Our discovery that MirA drives actin-based motility broadens the scope of host cellular processes targeted by PE_PGRS proteins. Moreover, we identify the specific host target of MirA, the host protein N-WASP, and show that the MirA PGRS domain directly binds and activates N-WASP to promote actin polymerization.

MirA’s glycine-rich nature makes it unique among the known WASP/N-WASP-binding partners. At resting state, N-WASP activity is autoinhibited by intramolecular interactions between the regulatory GTPase binding domain (GBD) and the WH2-central-acidic (WCA) domain that binds to actin and the Arp2/3 complex (*39–41*). The three previously identified microbial proteins that bind directly to N-WASP (*Shigella flexneri* IcsA, enterohemorrhagic *Escherichia coli* EspFU, and *Chlamydia trachomatis* TmeA) target the N-WASP GBD (*42–46*). However, the GBD of WASP/N-WASP is dispensable for *M. marinum* actin-based motility, and instead MirA targets a small positively charged basic region only known to interact with a phosphoinositol lipid (PIP2) (*9, 47, 48*). The discovery of MirA provides a new model to understand how WASP family proteins can be regulated and could lead to the identification of glycine-rich eukaryotic proteins that regulate actin dynamics. Moreover, the MirA-N-WASP interaction represents a tractable model to understand how other PE_PGRS proteins intercept host cell biological pathways to promote mycobacterial pathogenesis.

## Materials and methods

### Bacterial strains, media, and plasmids

*Escherichia coli* strains XL1-blue and BL21(DE3) were obtained from the UC Berkeley MacroLab and were used for cloning and protein purification, respectively. *E. coli* was cultured in lysogeny broth (LB), and antibiotics were used at concentrations of 100 µg/ml ampicillin, 50 µg/ml zeocin, or 50 µg/ml kanamycin when appropriate. Standard techniques were employed for cloning and other genetic manipulations.

*M. marinum* strain M (NCBI:txid216594) (*51*) was cultured in Middlebrook 7H9 broth (Fluka, M0178) supplemented with 0.2% glycerol, 0.05% Tween 80, and 10% oleic acid-albumin-dextrose-catalase (OADC), or on 7H10 agar (Difco, 262710) plates supplemented with 0.2% glycerol and 10% OADC. When appropriate, antibiotics were added to the media at the following concentrations: 50 μg/ml hygromycin, 20 μg/ml kanamycin, and/or 30 μg/ml zeocin. *M. marinum* was grown at 33°C and with shaking at ∼100 rpm for liquid cultures. Electroporations to introduce DNA were conducted as previously described(*52*).

### Cell lines

Mammalian cell lines U2OS (RRID:CVCL_0042), Raw 264.7 (RRID:CVCL_0493), HEK293 (RRID:CVCL_0045), and A549 (RRID:CVCL_0023) were obtained from and authenticated using short-tandem-repeat analysis by the University of California, Berkeley Cell Culture Facility (UCB-CCF). Cell lines were confirmed to be mycoplasma negative by DAPI staining and fluorescence microscopy screening at the UCB-CCF. Cells were grown at 37°C in 5% CO2 and maintained in DMEM (Gibco; 11965-092) containing 2-10% heat-inactivated and filtered fetal bovine serum (FBS) (Atlas Biologicals, FP-0500- A). Sf9 insect cells (RRID: CVCL_0549) were also obtained from the UCB-CCF and grown at 28°C in in Grace’s insect media (Gemini Bio-Products, 600-310) supplemented with 2% FBS (Gemini Bio-Products, 100-500) and penicillin/streptomycin. Bone marrow-derived macrophages from wild-type C57BL/6J mice were isolated and derived exactly as previously detailed(*53*). Polyclonal U2OS cells stably expressing F-Tractin-mCherry (*54*) to mark F-actin or A549 cells expressing a farnesylated TagRFP to mark the plasma membrane were generated by lentiviral transduction as described previously (*55*).

### Transient transfections

Cells were plated in 24-well plates 18-24 h prior to transfections at 1-1.5 x 10^5^ cells/well such that cells were 60-80% confluent. Plasmid DNA (500 ng) was diluted in Opti-MEM (Thermo, 31985062) and mixed with Lipofectamine 2000 (Thermo, 1668019) as per manufactures protocol. Cells were washed once with phosphate-buffered saline (PBS) (Gibco, 10010-023) and incubated in 300 µl Opti-MEM. DNA-Lipofectamine 2000 complexes were added to the cells for 2-4 h at 37°C, then replaced with DMEM + 2% FBS. Samples were fixed or lysed at 24-48 h post transfection. For transfections coupled to bacterial infection, cells were initially transfected for 4 h then infected with *M. marinum* and harvested 48 hpi.

For eukaryotic expression of *mirA*: wild-type (pBH261; *M. marinum M* (NCBI Reference Sequence: NC_010612.1) nts 4,401,156-4,403,237), ΔPE (pBH310; nts 4,401,447-4,403,237), and ΔAH (pBH351; nts 4,401,156-4,401,453 and 4,401,519-4,403,237) versions were cloned into the NotI/BsrGI site of pBH257 with 5’ ATG and consensus Kozak sequence with a 3’ V5 and 6xHis epitope tags. To create the *mirA* amphipathic helix fused to tdTomato (pBH347), nts 4,401,447-4,401,549 from *M. marinum* genomic DNA was amplified and fused upstream to *tdTomato* amplified from pBH251 as template, then Gibson cloned into the NotI/BsrGI site of pBH257 with a 5’ ATG and consensus Kozak sequence. In **Figure 3G**, U20S cells were transfected with pBH347, fixed at 24 hpt, then stained with BODIPY 493/503. For **Figures 3A-D**, cells were transfected with a plasmid encoding the empty parental vector (pBH257), *PEF-1α*-*mirA*^261-2049^*-V5* (pBH310), or *PEF-1α*-*mirA*^Δ297-363^*-V5* (pBH351). For **Figure S3D**, MMAR_2645 (nts 3,221,630-3,223,507) was cloned into the NotI/BsrGI site of pBH257 with 5’ ATG and consensus Kozak sequence and a V5 epitope tag creating pBH443. Cells were fixed 24 h post-transfected in U2OS cells and stained with for MMAR_2645 (*α*-V5) and lipid droplets (BODIPY 493/503). For **Figure 4A**, U20S cells were transfected with either the empty parental vector (pBH257) or *PEF-1α*-*mirA^ΔPE^-V5* (pBH310). At 24 hpt, cells were fixed and stained for lipid droplets (BODIPY 493/503), MirA (*α*-MirA), and either N-WASP (*α*-N- WASP) or the Arp2/3 complex (*α*-P34).

### Bacterial infections of host cells

Host cells used for infection were seeded ∼18-24 h prior to infection. U2OS cells were seeded into 96- and or 24-well plates at 4 x 10^4^ or 2 x 10^5^ cells per well, respectively. U2OS cells were incubated in DMEM without serum for ∼60 min prior to infection to promote uptake of *M. marinum*. RAW 264.7 cells were seeded at 7.8 x 10^6^ into T-75 flasks. BMDM were seeded into 96- or 24-well plates with 5 x 10^4^ and 3 x 10^5^ macrophages per well, respectively.

Bacteria were recovered from −80°C and resuspended into ∼2-3 ml of 7H9-OADC at an OD600 ∼0.1-0.3. Cultures were grown for 18-24 h, then back-diluted into fresh 7H9-OADC to an OD600 ∼0.1 and grown for an additional ∼18-24 h to an OD600 ∼0.8 to standardize growth phase prior to infection. Prior to infections, *M. marinum* was washed 3 times in 1 ml PBS, assessed for OD600 to calculate MOI, then resuspended in DMEM + 10% serum. Bacteria were added to pre-seeded mammalian cells and immediately spinfected at 1000 rpm for 10 min at 22°C. Infections were moved to a 33°C incubator for 2 h before rinsing 3 times with PBS, then maintained in DMEM with 2-10% FBS with amikacin (40 µg/ml) to kill extracellular bacteria at 33°C.

To measure infectious focus size, U2OS cells were plated onto 12 mm coverslips in 24- well plates. 18-24 h later, confluent monolayers were infected at an MOI of 1-20. Infection progressed for indicated time at 33°C until fixation and staining. To quantify spread, individual foci were assessed for the number of infected host cells per focus where *M. marinum* was marked by expression of tdTomato from a multicopy plasmid (pBH251) and boundaries of host cells were determined by cortical actin staining via Alexa 488 phalloidin. For **Figures 1B and 1C**, WT (BHm136), Δ*mirA* (BHm132), or Δ*mirA* + *attB*::*PmirA*-*mirA* (BHm163) strains were used and >20 foci were counted per replicate over three biological replicates. For **Figure S1A**, WT (BHm102), *espG5*::*tn*-1 (BHm106), or *espG5*::*tn*-2 (BHm107) were used to infect U2OS cells and >12 foci were counted per replicate over three biological replicates. For **Figure S1I**, WT (BHm136), and Δ*mirA* + *PgroEL*-*3581* (BHm152) were used to infect U2OS cells and >20 foci were counted per replicate over three biological replicates.

To measure the colocalization of *M. marinum* associated with N-WASP, MirA, and/or F-actin, U2OS or BMDM cells were plated on 12 mm coverslips and infected 18 h later at an MOI 1-50. Infected cells were incubated at 33°C for the indicated time and fixed and stained for N-WASP (*α*-N-WASP), MirA (*α*-MirA), and/or F-actin (Alexa 488 phalloidin). For **Figures 1E and 1F**, WT (BHm103), Δ*mirA* (BHm144), and Δ*mirA* + *P3581*-*3581* (BHm161) were used and 50-400 bacteria/replicate were calculated over three biological replicates. For **Figures S1D and S1E**, WT (BHm103) and Δ*mirA* (BHm144) were used to infect C57BL/6J BMDM cells and fixed at 48 hpi and stained for F-actin (Alexa 568 phalloidin). >300 bacteria were assessed for both strains for an actin tail for three biological replicates. For **Figures S1F-I**, WT (BHm103), Δ*mirA* (BHm144), Δ*mirA* + *P3581*- *3581* (BHm170), and/or Δ*mirA* + *PgroEL*-*3581* (BHm149), were used in U2OS cells. For **Figure S1F**, 50-250 bacteria/replicate were assessed for colocalization with both F-actin (Alexa 488 phalloidin) and N-WASP (*α*-N-WASP) over three biological replicates. **Figure S1G** was fixed at 72 hpi and stained for F-actin (Alexa 488 phalloidin). To measure the bacterial cell length of *M. marinum* during infection in **Figure S1H**, 50-150 bacteria from each strain and time point were assessed for length using the freehand line tool in ImageJ (*56*) from three biological replicates.

To calculate bacterial growth during infection shown in **Figures 1D** and **S1C**, WT (BHm103) or Δ*mirA* (BHm144) bacteria were used to infect U2OS or BMDM host cells (in a 24-well plate format) at an MOI of 0.1 or 0.01, respectively. Prior to harvesting, infected cells were washed three times with PBS to remove extracellular bacteria. Host cells were then lysed in water for 10 min at 37°C. From those lysates, serial dilutions were made in 7H9-OADC, plated onto 7H10-OADC + hygromycin agar plates, then enumerated 8-10 d later. Data represents three biological replicates. To calculate bacterial growth in media (**Figure S1B**), WT (BHm101), Δ*3581* (BHm129), and Δ*mirA* + *P3581*-*3581* (BHm161) strains were cultured in 7H9-OADC broth at 33°C/100rpm and monitored by A600 over two biological replicates.

For examining MirA by immunofluorescence and colocalization with F-actin and N-WASP in **Figures 2B-D** and **3I**, Δ*mirA* (BHm144), Δ*mirA* + *PmirA*-*mirA-flag* (BHm170) or Δ*mirA* + *PmirA*-*mirA^ΔAH^-flag* (BHm174) 50-300 bacteria were assessed for MirA localization for each strain at every time point over three biological replicates. The infection shown in **Figure 3I** was fixed at 48 hpi and stained for the FLAG epitope and F-actin (Alexa 488 phalloidin). For **Figure 2E**, Δ*mirA* + *PmirA*-*mirA-flag* (BHm170) was assessed at each time point for colocalization with N-WASP (*α*-N-WASP) or an actin tail (Alexa 488 phalloidin) in U2OS cells counting 150-200 bacteria per time point over three biological replicates. In **Figures 2F and 2G**, Δ*mirA* + *PmirA*-*mirA-flag* (BHm170) was used to infect U2OS cells and 100-150 bacteria were assessed at each time point for three biological replicates. For examining *in trans mirA* complementation of Δ*mirA* bacteria, U20S cells were transfected with either a plasmid encoding *mirA-V5* (pBH261) (**Figure 2H**), *mirA^ΔPE^-V5* (pBH310) (**Figure S2E**), or *mirA^ΔAH^-V5* (pBH351) (**Figure 3H**). Transfections were washed and then infected with *M. marinum* Δ*mirA* (BHm132), then fixed at 48 hpi and stained for MirA-V5 (*α*-V5) and F-actin (Phalloidin-iFluor 405). For **Figure S2D**, 50-150 bacteria/replicate of each MirA variant were assessed for association with MirA or F-actin for three biological replicates.

### Generating, screening, and mapping a *M. marinum* transposon insertion library

A *M. marinum* transposon insertion library expressing mCherry was generated through transduction of the mycobacterium-specific phage phiMycoMarT7 containing the mariner-like transposon Himar1 as previously described (*57, 58*). This library had an estimated ∼4.4 million insertions (∼35-fold coverage of TA dinucleotide insertion sites in *M. marinum*). Individual colonies were isolated from 7H10-OADC plates containing kanamycin and hygromycin, grown in 80 µl of 7H9-OADC with containing kanamycin and hygromycin in 96-well plates, and used to infect near-confluent U2OS cells in a 96-well format. At 72 hpi, clones with defects in bacterial cell-to-cell spread were identified by fluorescence microscopy. Transposon insertions that caused defects in cell spread due to a growth attenuation (e.g., insertions into genes of the ESX-1 secretory system) were not further examined.

Semi-random nested PCR was used to map the transposon insertion site of spread- defective mutants. To map the 5’ direction, oligos GCTTAGTACGTTAGCCATGAGAGC and GGCCACGCGTCGACTAGTCANNNNNNNNNNAGCTG were used in PCR with chromosomal DNA as the template. Subsequently, those PCR products were used as template with oligos CACATTTCCCCGAAAAGTGCCAC and GGCCACGCGTCGACTAGTCA. To map the 3’ direction oligos TACCTGCCCATTCGACCACCAAGC and GGCCACGCGTCGACTAGTCANNNNNNNNNNAGCTG used in PCR with chromosomal DNA as the template. Subsequently, those PCR products were used as template with oligos CGCATCGCCTTCTATCGCCTTCTT and GGCCACGCGTCGACTAGTCA. ExoSAP-IT was then mixed with the second PCR and incubated for 30 min at 37°C, then heat inactivated. The amplicons were then sequenced with GGCCACGCGTCGACTAGTCA and genomic locations were determined using BLAST (GenBank/NCBI accession NZ_HG917972). A transposon insertion into *MMAR_3581* (*mirA*) was identified at nucleotide 4,403,232 (2046 in CDS) (strain BHm119). Two distinct transposon insertions into MMAR_2680 (*eccA5*) were identified, at nucleotide 3,269,477 (1561 in CDS) (strain BHm106) or 3,269,674 (1758 in CDS) (strain BHm107).

### Construction of the *mirA* deletion and complementation strains

The *M. marinum* Δ*mirA* strain was built using the *sacB*/*galK* counterselection method(*59*). Briefly, ∼1000 bp from both the 5’ or 3’ end of *mirA* was cloned into pBH92 resulting in pBH263. WT *M. marinum* was electroporated with pBH263 and plated on 7H10-OADC with 50 μg/ml hygromycin, then screened for recombination by PCR. Strains with the initial recombination were grown to an A600 of ∼1.0 in 7H9-OADC and plated onto 7H10-OADC containing 5% sucrose and 0.2% 2-deoxy-galactose to select strains that had undergone the second recombination. Strains with a clean *mirA* deletion were identified by PCR and DNA sequencing.

The Δ*mirA* strain was complemented with MirA at two different expression levels. First, an over-expression strain was built by amplifying full-length *mirA* from *M. marinum* genomic DNA (nts 4,401,186-4,403,237) and cloned into pMV261 (*60*) (pBH206) downstream of *PgroEL* to create pBH417. Either *PMSP12*-*ebfp2* or *PMSP12*-*tdtomato* was then cloned into the PciI site to create pBH423 or pBH424, respectively. Second, a native MirA expression strain was built by inserting the *mirA* locus (nts 4,398,526-4,403,237) into the HindIII/KpnI site of the integrative vector pST-Ki (*61*) (pBH255) thereby creating pBH445. The *mirA* amphipathic helix deletion was built similarly, however it was amplified in two parts thereby omitting nts 4,401,492-4,401,548 creating pBH450. pBH445 or pBH450 were electroporated into Δ*mirA* expressing EBFP2 (BHm144) creating the BHm170 or BHm174 *M. marinum* strains.

### Immunoblotting

For immunoblotting to evaluate protein expression, cells were lysed in ice-cold RIPA buffer (Thermo, 89900) with protease inhibitors, protein concentrations were measured, then samples were mixed with 4x sample buffer and immediately cycled 3 x 5 min between a 100°C water bath and ice. Lysates were electrophoresed by SDS-PAGE, then applied to PVDF membrane via semi-dry protein transfer. Membranes were then blocked in 5% milk, washed with PBS, then incubated with primary antibodies overnight at 4°C. Primary antibodies were diluted in blocking buffer (2% BSA and 0.1% Tween-20 in PBS) as follows: GroEL (1:1000), GAPDH (1:10,000), N-WASP (1:1000), CDC-42 (1:750), WIPF2 (1:750), MirA (1:7500), His (1:1000), and Strep (1:1000). Washed membranes were then incubated in 1% milk with 1:5000 secondary antibodies conjugated to HRP then imaged.

For immunoblotting of mycobacterial proteins, bacteria were isolated by centrifugation. Pellets were resuspended into PBS and heat inactivated for 60 min at >90°C. Bacteria were pelleted and resuspended in mycobacterial lysis buffer (15% sucrose, 50 mM Tris-pH 8.5, 50 mM EDTA), 1.0 mm silica beads were added, then lysed cells using a Mini Beadbeater-8 Cell Disrupter (Biospec Products) at maximum power for 3 x 40 s. Protein concentrations were measured, then 4x sample buffer was added and cycled 3 x 5 min between a >90°C water bath and ice. Lysates were then electrophoresed and immunoblotting as described above. For **Figures 2A and S2B**, expression levels were normalized to 72 h as 100% (n = 3), and WT (BHm103) and Δ*mirA* (BHm144) strains were used. For **Figure S2A**, MirA^WT^ (BHm144), MirA^+++^ (BHm149), MirA^WT^-Flag (BHm170), and MirA^ΔAH^-Flag (BHm174) were used with GroEL2 as the loading control. For **Figure S2C**, plasmids encoding either empty (pBH257), MirA^WT^ (pBH261), MirA^ΔPE^ (pBH310), or MirA^ΔAH^ (pBH351) were transfected in U2OS cells.

### Fluorescence microscopy

Eukaryotic cells affixed to 12 mm glass coverslips in 24-well plates were fixed with 4% paraformaldehyde in PBS for 10 min at 22°C. Cells were then permeabilized with 0.5% Triton X-100 for 5 min, washed, then blocked with blocking buffer (2% BSA and 0.1% Tween-20 in PBS) for 20 min at 22°C. Primary antibodies were incubated with samples overnight at 4°C in a humidified chamber. Primary antibodies were diluted in blocking buffer as follows: N-WASP (1:750), CDC-42 (1:1000), Arp2/3 (1:500), MirA (1:1000), V5 (1:1000) and FLAG (1:1000). Secondary antibodies, conjugated to Alexa fluorophores, were used at 1:1000 in blocking buffer. F-actin was stained using Alexa 568 phalloidin (Thermo, B3475) or Alexa 488 phalloidin (Thermo, A12379) at 1:400, or Phalloidin-iFluor 405 (Abcam, ab176752) at 1:250, in blocking buffer. For experiments staining lipid droplets, cells were permeabilized, blocked, and maintained in 2% BSA with 0.5% saponin in PBS. BODIPY 493/503 (Thermo, D3922) was added at a concentration of 1 µM, and coverslips were mounted in 20 mM Tris, 0.5% N-propyl gallate, 85% glycerol, pH 8. Images were captured on a Nikon Ti Eclipse microscope with a Yokogawa CSU-XI spinning disc confocal, 60X and 100X (1.4 NA) Plan Apo objectives, a Clara Interline CCD Camera and MetaMorph software. Images were processed and quantified using ImageJ and assembled in Adobe Illustrator.

### Movement parameters of *M. marinum* and lipid droplets

The velocities of *M. marinum* and lipid droplets were measured using U2OS cells stably expressing F-tractin-mCherry. Cells were seeded at 6 x 10^5^ in 20mm MatTek glass bottom dishes, incubated for 18-24h then either infected with *M. marinum* expressing GFP or transfected with a plasmid encoding with *mirA*^ΔPE^*-V5* (pBH310). Bacteria were evaluated 48-72 hpi, while lipid droplets were imaged 24-48 hpt. Live-cell imaging was conducted in a 33°C environmental chamber with a Nikon Ti Eclipse with 100x objective at 5” intervals for 5-10’. Only lipid droplets with a diameter similar to the width of mycobacteria (0.35 > x > 0.75 µm) were evaluated. To calculate actin-tail mediated velocities, the manual tracking plugin in ImageJ was used to track movement >10 consecutive frames (52 tracks were collected for *M. marinum* and 88 tracks for lipid droplets over four experiments). The efficiency of movement was measured by calculating final displacement divided by the total distance traveled over a 100” time interval (n = 78). The actin tail length was measured from still frames of live-cell microscopy (n = 89).

### Lipid droplet cell-to-cell spread

For monitoring lipid droplet cell-to-cell spread, A549 cells stably expressing farnesylated- RFP to mark the plasma membrane (donor cells) were plated in a 24-well plates. 18-24 h after plating, donor cells were transfected with pBH310 (*PEF-1a*-*mirA^ΔPE^*). At 4 hpt, donor cells washed 3x PBS, lifted with 0.25% trypsin. Donor cells were then mixed with unmarked, untransfected A549 (recipient cells) at a ratio of 1:10 and plated on glass coverslips in a 24-well plate. Plates were placed in a humidified secondary container in the incubator to promote even cell distribution. Cells were incubated for 36 h at 37°C prior to fixation and staining.

### Affinity purification and mass spectrometry

Affinity purification and mass spectrometry was preformed similar to previously described (*62*)*. mirA* without the PE super domain (nts 4,401,447-4,403,237) was amplified by PCR from *M. marinum* genomic DNA with a C-terminal twin strep tag and cloned into the NotI/BsrGI site of pBH257 resulting in pBH342. Additionally, the amphipathic helix mutant was built similarly except mirA^ΔAH^ was generated from PCR with pBH351 as template to create pBH364. pBH342 or pBH364 and the parental vector (pBH257) were separately transfected into HEK293 cells in 4 x 8 cm plates using calcium phosphate-mediated transfection. At 42 hpt, cells were scraped, pelleted for 5’ at 1000 rpm and 4°C, then resuspended in cold lysis buffer (50 mM Tris-HCl, 150 mM NaCl, 1 mM EDTA, 0.5 % IGEPAL CA-630, protease inhibitors, pH 7.4) and lysed using a Dounce homogenizer. Lysates were clarified by centrifuging for 20’ at 2800 g and 4°C. Supernatants were initially incubated with 120 µl preclearing beads (Sepharose FF) for 2 h with rotation at 4°C. The precleared lysates were immobilized with Strep-Tactin resin (15 µl/8 cm plate) for 4°C for 18 h with rotation. The resin was washed 5x lysis buffer, and twice in lysis buffer without detergent before elution in that buffer containing 2.5 mM desthiobiotin (40 µl/8 cm plate). The eluate was analyzed by SDS-PAGE and either Coomassie or silver staining prior preceding with trypsin digestion. To use N-WASP as bait, full-length N-WASP was amplified from pBH5 adding a 5’ twin strep tag using an oligonucleotide primer. Strep-N-WASP was inserted into the XbaI site (under *PCMV*) of pBH310 that co-expressed *mirA^ΔPE^-V5* creating pBH359. Experimental conditions were as described above, then eluates were probed for N-WASP and MirA via immunoblot. For **Figures S4A and S4B**, HEK293 cells were transfected with a plasmid encoding either the empty parental vector (pBH257) or *mirA*^ΔPE^*-strep-his* (pBH342). For **Figure S4C**, HEK293 cells were transfected with a plasmid encoding either *mirA^ΔPE^-V5* (pBH310) or *mirA^ΔPE^-V5* and *strep-n-wasp* (pBH359). For **Figure S4D**, HEK293 cells were transfected with a plasmid encoding either the empty parental vector (pBH257), *mirA*^ΔPE^*-strep-his* (pBH342), or *mirA*^ΔPE,Δ*AH*^*-strep-his* (pBH364).

For subsequent mass spectrometry, proteins eluates were digested with trypsin for LC-MS/MS analysis. Samples were denatured and reduced in 2 M urea, 10 mM ammonium bicarbonate, 2 mM DTT at 60°C for 25 min then immediately alkylated with 2 mM iodoacetamide for 30 min in the dark at 22°C. Trypsin was added at a 1:100 trypsin: substrate ratio and digested overnight at 37°C. Samples were desalted using C18 ZipTips (Millipore) then speed vacuumed to desiccation and finally resuspended in 0.1% formic acid.

Digested peptide mixtures were analyzed by LC-MS/MS on a Synapt G2-Si mass spectrometer, which was equipped with a nano electrospray ionization source (Waters). The mass spectrometer was connected in line with an Acquity M-class ultra-performance liquid chromatography system equipped with trapping (Symmetry C18; inner diameter, 180 μm; length, 20 mm and particle size, 5 μm) and analytical (HSS T3; inner diameter, 75 μm; length, 250 mm; particle size, 1.8 μm) columns (Waters). Data-independent, ion mobility-enabled, high-definition mass spectra and tandem mass spectra were acquired in the positive ion mode. Data acquisition was controlled using MassLynx software (version 4.1), and tryptic peptide identification and relative quantification using a label-free approach were performed using Progenesis QI for Proteomics software (version 4.0, Waters). The raw data were matched to protein sequences by the Protein Prospector algorithm (*63*). Data were searched against a database containing SWISS-PROT Human protein sequences and concatenated to a decoy database where each sequence was randomized to estimate the false positive rate (accessed 28 August 2018). Protein species with a normalized spectral abundance factor of over 0.00025 were then scored with <u>M</u>ass spectrometry Interaction <u>ST</u>atistics (MiST) algorithm, using the MiST reproducibility (0.45), specificity (0.50) and abundance (0.05) weights (*23*). MirA-prey pairs with a MiST score ≥ 0.75 were deemed confident interactions and were combined with human protein interactions from the STRING databases (*49*). The resulting network diagram from two biological replicates was created using Adobe Illustrator in **Figure 4B**.

### Protein Purification

To generate MBP-fused MirA for initial affinity purification, *mirA* (pBH312; nts 4,401,156- 4,403,237) or *mirA^Δ1-297^* (pBH324; nts 4,401,447-4,403,237) were PCR amplified from *M. marinum* genomic DNA and cloned in-frame into the SspI site of 6XHis-MBP-TEV (AddGene: 29656). Plasmids were freshly transformed into the *E. coli* BL21(DE3) background and MirA expression was induced at 37°C with 0.5 mM IPTG for 3 h. His-MBP-MirA were first isolated using amylose bead affinity chromatography (NEB) in conditions specified by the manufacturer. His-MBP was cleaved from MirA overnight at 4°C in the presence of the His-TEV at a molar concentration of 1:100. MirA was purified away from His-MBP by gel-filtration chromatography using a Superdex 200 column in 20 mM Tris-HCl (pH 7.5), 300 mM NaCl, 10% glycerol, 0.5 mM DTT, then snap frozen in liquid nitrogen.

His-N-WASP was expressed and purified using the baculovirus expression system. His- N-WASP was PCR amplified from pBH5 using primers encoding a 6x-his and cloned into a bacmid resulting in pBH354. Recombinant baculovirus was generated in Sf9 cells using the Bac-to-Bac system according to the manufacturer’s instructions (Invitrogen, Carlsbad, CA). Recombinant proteins were then expressed by infecting Hi5 cells for 96 h at 28°C. Hi5 cells were pelleted, resuspended in lysis buffer (20 mM NaH2PO4, 400 mM NaCl, 20 mM imidazole, 10 mM BME, 1% IGEPAL, 25 units/l benzoate nuclease, protease inhibitors, pH 7.8), incubated for 15’ on ice, then sonicated at 10% power 4 x 15”. Lysates were spun at 20K rpm at 4°C for 45’. His-N-WASP was first isolated using Ni-NTA beads (Qiagen, Valencia, CA) using a buffer with an imidazole gradient (20 mM Bis-Tris, 500 mM NaCl, 1 mM TCEP, pH 6.5, imidazole 20 mM -> 500 mM). Fractions were then concentrated and flowed through a 5 mL SP SH cation exchange column to a cation exchange column (20 mM Bis-Tris, 100 -> 1000 mM KCl, 2 mM MgCl2, 0.5 mM EGTA, 0.5 mM EDTA, pH 6.5). Fractions were then subjected to gel filtration using a Superdex 200 column into Control Buffer (20 mM Bis-Tris, 400 mM KCl, 2 mM MgCl2, 0.5 mM EGTA, 0.5 mM EDTA, 0.5 mM DTT, 10% glycerol, pH 6.5). Fractions were concentrated and snap frozen in liquid nitrogen.

### MirA antibody

Polyclonal antibodies were generated against MirA in a rabbit host. The native full-length MirA protein was purified as described above and sent to Pocono Rabbit Farm and Laboratory (Canadensis, PA) where a 91-day custom antibody protocol was performed for both organisms. To affinity purify MirA specific antibodies from serum, purified MirA^ΔPE^ (described above) was dialyzed into ligand coupling buffer (200 mM NaHCO3, 500 mM NaCl, pH 8.3) and then coupled onto NHS-ester Sepharose 4 Fast Flow resin. The resin was incubated with serum overnight at 4°C with rotation, washed, and eluted with low pH buffer (100 mM Glycine, pH 2.5). Eluted fractions were neutralized with 1 M Tris pH 8.8 and dialyzed into PBS overnight at 4°C. Affinity purified antibodies were then concentrated using an amicon filter and stored at −80°C.

### Bulk Actin Assembly Assays

Rabbit skeletal muscle actin was purchased (unlabeled and pyrene-labeled actin; Cytoskeleton Inc.). All actin was maintained in G buffer (5 mM Tris, pH 8.0, 0.2 mM CaCl2, 0.2 mM ATP, 0.5 mM DTT) prior to polymerization assays. Pyrene actin polymerization reactions were started by combining monomeric actin in G buffer (at a final concentration of 1 μM actin (10% pyrene-labeled) with a mix of MirA, 10X initiation buffer (10 mM MgCl2, 10 mM EGTA, 5 mM ATP, 500 mM KCl) and MirA buffer (see purification details). Fluorescence was detected at 20 s intervals for 1 h on a Tecan Infinite F200 Pro plate reader using a 365 nm excitation filter (10 nm bandpass), a 405 nm emission filter (20 nm bandpass) and Magellan software. The reported reaction curves were normalized to the minimum fluorescence value of each reaction. The time for each reaction to reach the half-maximum fluorescence was determined following normalization to both the minimum and maximum fluorescence values, averaged from a minimum of three experiments as previously described (*64*). The means are reported with their standard deviations. Data were analyzed in Excel, and the final graphs were constructed in Prism v8.0 (GraphPad Software, Inc). For **Figures 4F and 4G**, 1 µM actin (10% pyrene-labeled) was mixed with initiation buffer alone or with 200 nM MirA, 200 nM MirA + 50 nM Arp2/3, 200 nM MirA + 200 nM N-WASP. Data in **Figure 4G** is mean ± SD from 3 technical replicates. For **Figures 4H and 4I**, 1 µM actin (10% pyrene-labeled) was mixed with either 50 nM Arp2/3, 50 nM Arp2/3 and 200 nM N-WASP, or 50 nM Arp2/3, 200 nM N-WASP and increasing concentrations of MirA. Data in **Figure 4I** is mean ± SD from 3 technical replicates.

### Pulldown assay

Purified native MirA (prey) and 6xHis-N-WASP (bait) were purified as described above. 50 µg of His-N-WASP was incubated alone or with 50 µg of MirA in 400 µL of for 2 h at 4°C under agitation. His-N-WASP ± MirA was loaded onto a column with 80 µL of previously equilibrated Ni-NTA resin (Qiagen) and incubated for 1 h at 4°C with agitation. Column was centrifuged for 1’ at 1000 rpm. Beads were washed thrice with 20 mM Tris– HCl, pH 7.5, 250 mM NaCl, 20 mM imidazole, then eluded with 100 µl of the same buffer except with 500 mM imidazole. Eluates were evaluated by immunoblot because MirA and N-WASP have equivalent molecular weights.

### Actin polymerization on beads in *Xenopus laevis* extract

Polystyrene microspheres (Fluoresbrite^TM^ Carboxylate YG, 0.5 μm and 1 μm, (Polysciences)) were incubated on ice with 5 μM MirA for >1 h before adding BSA to a concentration of 5 mg/mL and incubating for 15 min. Beads were washed in CSF-XB (10 mM HEPES at pH 7.7, 2 mM MgCl2, 0.1 mM CaCl2, 100 mM KCl, 5 mM EGTA and 50 mM sucrose) and kept at 4°C. *Xenopus laevis* egg extract was provided by the laboratory of Dr. R. Heald (University of California, Berkeley). To 8 μl of extract, 1 μl of actin (3 μM, 20% of which was rhodamine-labelled) and 1 μl of BSA- or MirA-coated beads were added, and 1-2 μl of this dispersion was placed between a slide and coverslip and observed immediately by epifluorescence microscopy. ImageJ was used to adjust brightness and contrast and export QuickTime movies. To calculate bead velocities, the manual tracking plugin in ImageJ was used to track movement 10 consecutive frames (every 2.5 s for the 0.5 µm beads and 5 s for the 1 µm beads). >40 counts over three replicates were assessed and shown in **Figure 4D**.

### Transmission electron microscopy

For examining *M. marinum* escape from the mycobacterial-containing vacuole by electron microscopy, *M. marinum* wild type (BHm102) and *mirA*::*tn* (BHm119) strains expressing mCherry were used to infect U2OS host cells (MOI = 10) in 6-well plates. At 60 hpi, cells were scraped, pelleted at for 5’ at 1000 rpm at 22°C, then gently resuspended in fixative (1.5% paraformaldehyde, 2.0% glutaraldehyde and 0.03% CaCl2 in 0.05 M cacodylate buffer, pH 7.2). After 45’ at 22°C, the samples were embedded in 2.0% glutaraldehyde and 0.1 M cacodylate buffer, and 2% low-melting agarose and incubated overnight at 4°C. The samples were post-fixed the following day in 1% osmium tetraoxide and 1.6% potassium ferricyanide, dehydrated in a graded series of ethanol concentrations and embedded in EPON 812 resin prepared as follows: 11.75 g eponate 12, 6.25 g dodecenyl succinic anhydride and 7 g NADIC methyl anhydride were mixed before the addition of ml of the embedding accelerator benzyldimethylamine. Next, the samples were stained with 2% uranyl acetate and lead citrate. Images were obtained on a FEI Tecnai 12 transmission electron microscope and processed on ImageJ. To quantify *M. marinum* localized within a vacuole shown in **Figures 1G and 1H**, two biological replicates of wild type (BHm102) and *mirA*::*tn* (BHm119) were assessed with >65 bacteria per replicate.

### Quantification and statistical analysis

Statistical parameters and significance are reported in the figures and the figure legends. Data are determined to be statistically significant when p < 0.05 where indicated. As such, asterisks denote statistical significance as: *, p < 0.05; **, p < 0.01; ***, p < 0.001; ****, p < 0.0001, compared to indicated controls. All other graphical representations are described in the figure legends. Statistical analysis was performed in GraphPad PRISM 9.

## Acknowledgements

We thank the laboratories of Sarah Stanley (UC Berkeley), David Tobin (Duke Univ.), James Olzmann (UC Berkeley), Jeffery Cox (UC Berkeley), and Christina Stallings (Washington Univ.) for providing us with reagents, equipment, and expertise. We thank Erin Benanti for purification of the Arp2/3 complex and generating the F-tractin-mCherry U20S cell line, Taro Ohkawa for expertise in insect protein purification, Rebecca Lamason for generating the mCherry-CAAX A549 cell line, Thomas Burke and Patrik Engström for generating BMDM cells, Daniel Portnoy and Neil Fischer for their critical reading of this manuscript, and current and former Welch lab members for their invaluable contributions. We also thank UC Berkeley core facilities including: Ann Fisher and Alison Killilea (Cell Culture Facility), Reena Zalpuri (Electron Microscopy Facility), Lori Kohlstaedt (Vincent J. Proteomics/Mass Spectrometry Laboratory), and Mary West (CTAF/HTSF). N.S.H. is supported by a Jane Coffin Childs Fund Postdoctoral Fellowship and M.D.W is supported by a National Institutes of Health grant R35 GM127108.

## Author Contributions

Conceptualization, NSH, MDW; Methodology, NSH, MDW; Investigation, NSH; Visualization, NSH; Funding acquisition, NSH, MDW; Project administration, NSH, MDW; Supervision, MDW; Writing – original draft, NSH; Writing – review & editing, NSH, MDW.

## Data Statement

All data are available in the main text or the supplementary materials.

---

**Figure S1.**
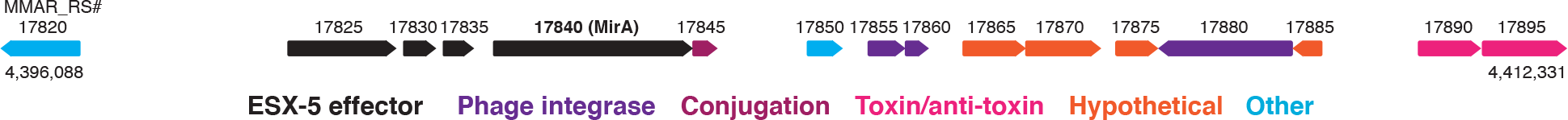
*mirA* locus is adjacent to factors associated with horizontal gene transfer. The *mirA* genetic locus (MMAR_RS17825 to MMAR_RS17840) is adjacent to elements associated with horizontal gene transfer, including genes coding for functions involved in conjugation, mycobacteriophage phage transduction, or a toxin/antitoxin system. Additionally, the locus is flanked by increased intergenic space (2173 nucleotides on the 5’ end and 991 nucleotides on the 3’ end), whereas the average intergenic space in the *M. marinum* genome is only ∼124 nt.

**Figure S2.**
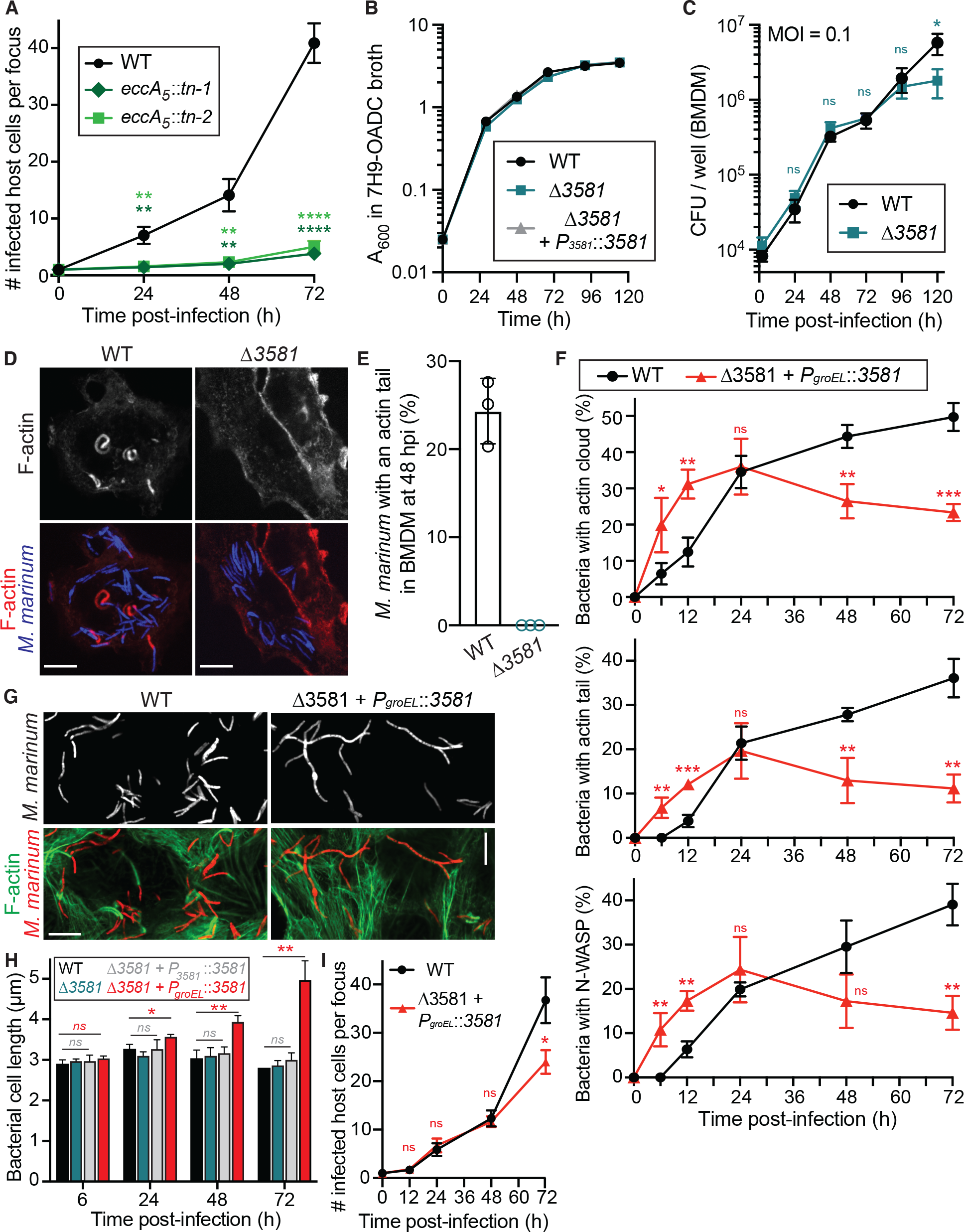
Growth dynamics of *mirA* knockout and overexpression. (**A**) Time course graph indicating the number of host U2OS cells in an infectious focus during infection of WT, *espG5*::*tn*-1, or *espG5*::*tn*-2. Data is mean ± SD; n = 3 (unpaired t-test). (**B**) Growth curve of WT, Δ*3581*, and Δ*3581* + *P3581*-*3581* strains in 7H9-OADC broth at 33°C. (**C**) Growth curve of WT versus the Δ*mirA* strains in BMDM host cells. Data is mean ± SD; n = 3. (**D**) Representative micrographs showing the lack of actin tail formation in Δ*mirA* bacteria in BMDMs at 48 hpi. Scale bar is 5 µm. (**E**) Comparing the number of actin tails between WT and Δ*mirA* bacteria in BMDM host cells, which express both nucleation promoting factors WASP and N-WASP. Data is mean ± SD; n = 3. (**F**) Time course graphs of *M. marinum* WT or MMAR_3581 overexpression strains colocalized with F-actin, an actin tails, or N-WASP. Data is mean ± SD (unpaired t-test); n = 3. (**G**) Representative micrographs of either *M. marinum* WT or MMAR_3581 overexpression strains during infection of U2OS cells at 72 hpi. (**H**) Cell length measurements of *M. marinum* WT, Δ*3581*, Δ*3581* + *P3581*-*3581*, Δ*3581* + *PgroEL*-*3581* over the course of an infection of U2OS cells. Data is mean ± SD (unpaired t-test); n = 3. (**I**) Time course graph indicating the number of host U2OS cells per infectious focus during infection of WT *M. marinum* WT or the Δ*3581* + *PgroEL*-*3581* overexpression strain. Data is mean ± SD (unpaired t-test); n = 3.

**Figure S3.**
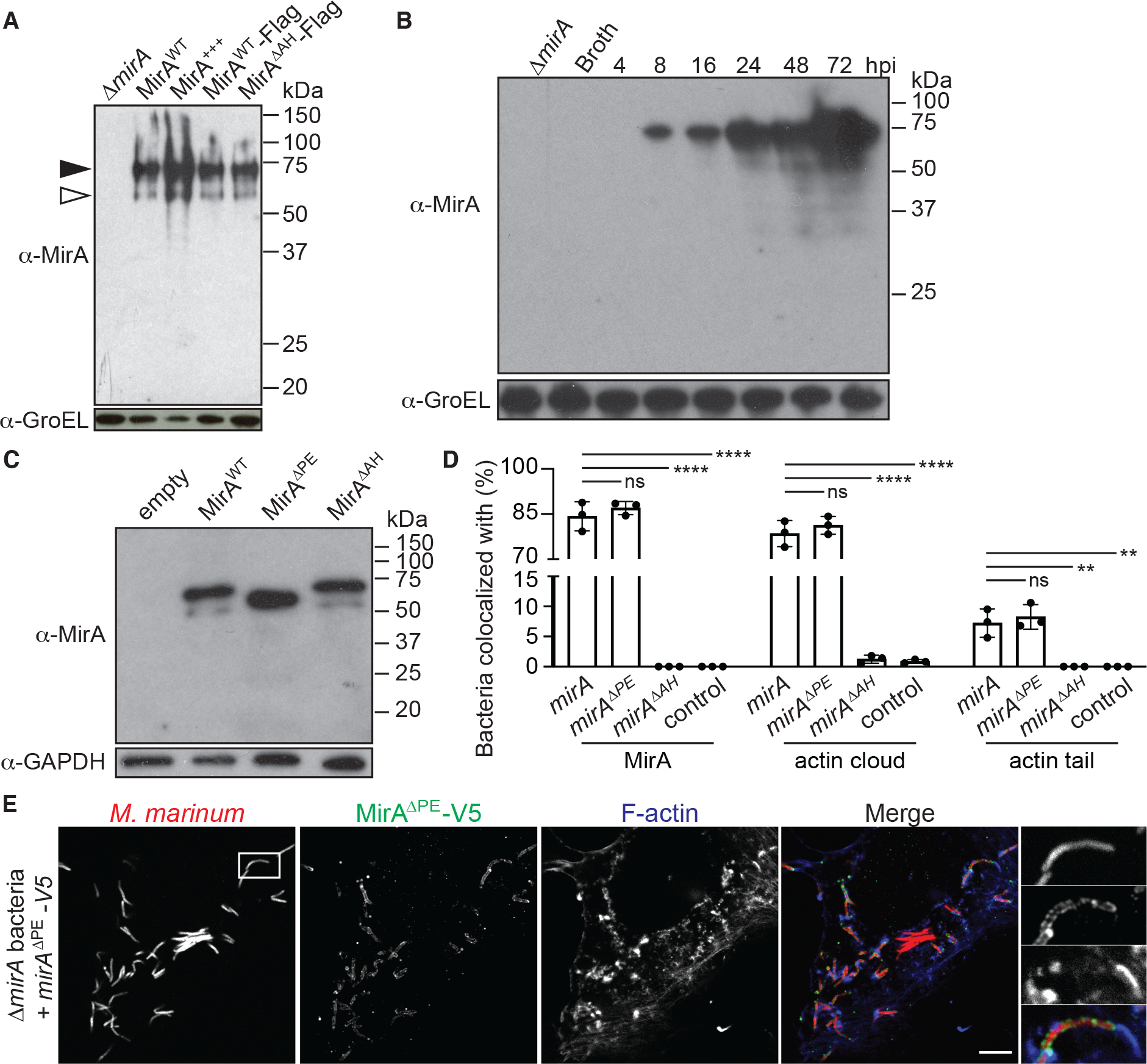
Endogenous and ectopic MirA expression levels and colocalization measurements for *mirA* complementation *in trans*. (**A**) Expression of MirA variants during infection of U2OS cells at 48 hpi. Closed arrow represents full-length MirA, while the open arrow likely represents a proteolytically processed MirA^ΔPE^ outer membrane form. GroEL expression is used as the loading control. (**B**) MirA expression during time course of infected RAW 264.7 macrophage cell line. GroEL is used as the loading control. (Image is the full version of Figure 2A.) (**C**) A representative immunoblot of ectopically expressing MirA variants in U2OS cells. GAPDH is used as the loading control. (**D**) Quantification of the percent of Δ*mirA M. marinum* with either MirA, actin clouds, or actin comet tail when supplied with ectopically expressed MirA variants. Data is mean ± SD; n = 3. (**E**) Ectopically expressed MirA^ΔPE^-V5 localizes to the surface of Δ*mirA* bacteria to stimulate actin polymerization at 48 hpi. White box indicates zoom panel and scale bar is 3 µm.

**Figure S4.**
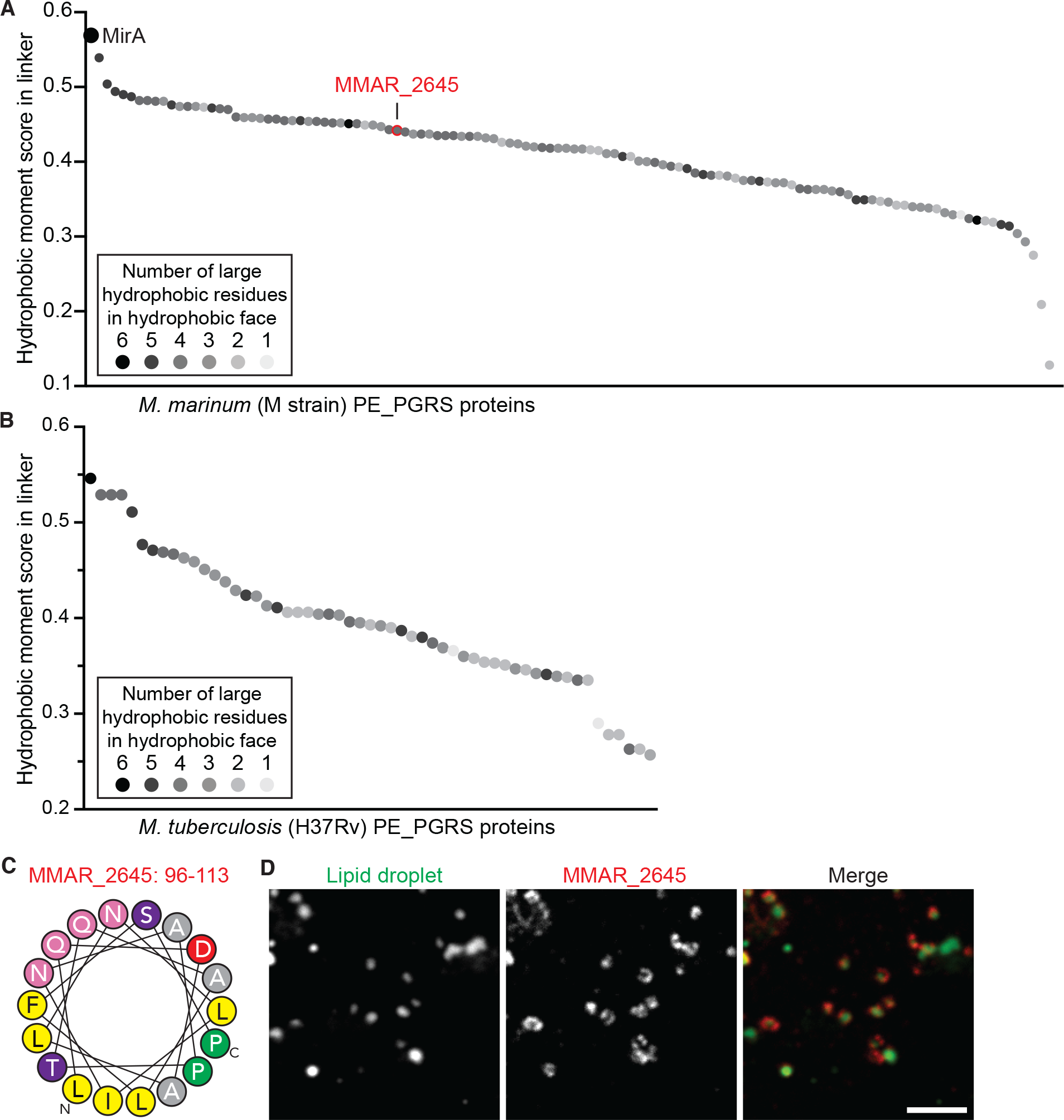
Detection of putative amphipathic helices within *M. marinum* and *M. tuberculosis* PE_PGRS proteins. (**A**) *M. marinum* PE_PGRS proteins were assessed for a putative amphipathic helix in the linker region between their PE and PGRS domains. The highest hydrophobic moment score is shown. Additionally, the number of large hydrophobic residues (I, F, L, M, W, Y) within the hydrophobic face, a predictive factor of amphipathic helix insertion into phospholipid monolayers (*50*), is shown in greyscale (inset). Further details are provided in **Data S1**. (**B**) PE_PGRS proteins from *M. tuberculosis* strain H37Rv were assessed for a putative amphipathic helix as described above and further detailed in **Data S1**. (**C**) The putative amphipathic helix in MMAR_2645 residues 96-114). (**D**) Full-length MMAR_2645-V5, encoding a more representative amphipathic helix hydrophobic moment score, was ectopically expressed in U20S cells and localizes to the surface of lipid droplet organelles similar to MirA. Scale bar is 2 µm.

**Figure S5.**
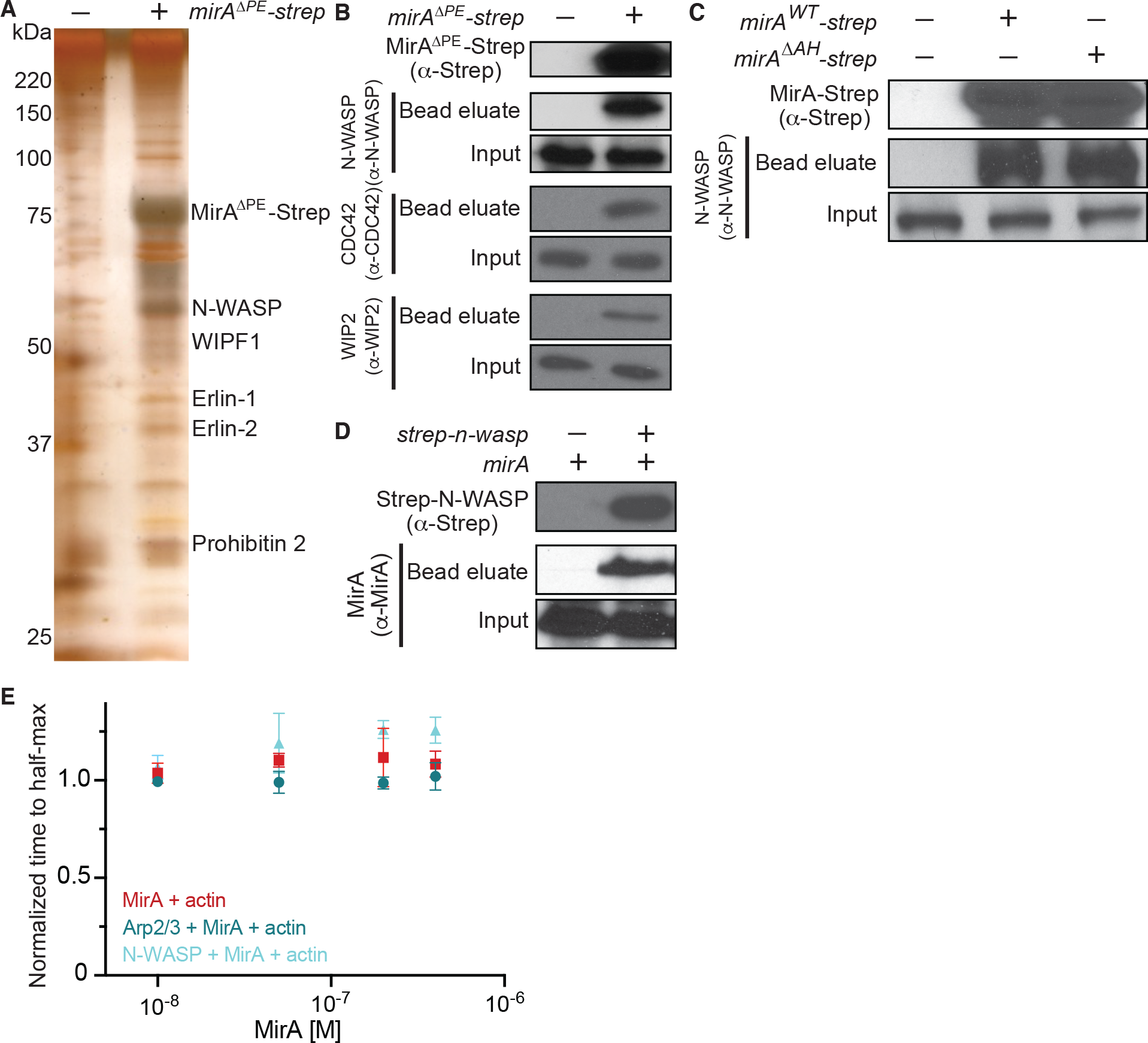
MirA interacts with N-WASP and other actin cytoskeletal proteins. (**A**) Eluates of Streptactin affinity purifications from HEK293 cells subjected SDS-PAGE and sliver stained. Proteins within the eluate identified by gel excision-mass spectrometry are indicated. (**B**) Representative immunoblots of MirA-Strep affinity purified lysates from HEK293 blotted against actin cytoskeletal proteins (N-WASP, CDC42, and WIP2) identified as MirA interactors by AP-MS shown in Figure 4B. (**C**) Representative immunoblots of the MirA-Strep amphipathic helix mutant affinity purified from HEK293 lysates against N-WASP. (**D**) Representative immunoblots of the Strep-N-WASP affinity purified lysates from HEK293 cells against MirA. (**E**) The time to half-maximum fluorescence of the polymerization curves normalized to actin alone. Data is mean ± SD; n = 3. Related to Figure 4F.

**Movie S1. Live-cell imaging of MirA-transfected cells that exhibit actin rocketing lipid droplets, Related to** Figure 3B. U2OS cells stably expressing a F-actin marker (F-tractin-mCherry) were transfected with *PEF-1α*-*mirA* (pBH261). Images were captured every 5 s for 10 m displayed at 8 frames/s and the movie is repeated twice. Timestamp shows min: sec and scale bar is 3 µm.

**Movie S2. MirA-transfected cells with labeled lipid droplets, Related to** Figure 3. U2OS cells stably expressing a F-actin marker (F-tractin-mCherry; red) were transfected with *PEF-1α*-*mirA^ΔPE^* (pBH310) and stained for lipid droplets (Bodipy 493/503; green). The F-actin channel alone is shown below. Images were captured every 5 s for 135 s displayed at 6 frames/s and movie is repeated three times. Timestamp shows min: sec and scale bar is 5 µm.

## References

1. C. M. Haglund, M. D. Welch, Pathogens and polymers: microbe-host interactions illuminate the cytoskeleton. J Cell Biol. 195, 7–17 (2011).

2. M. D. Welch, M. Way, Arp2/3-mediated actin-based motility: a tail of pathogen abuse. Cell Host Microbe. 14, 242–255 (2013).

3. J. N. Bugalhão, L. J. Mota, I. S. Franco, Bacterial nucleators: actin’ on actin. Pathog Dis. 73, ftv078 (2015).

4. S. Kühn, J. Enninga, The actin comet guides the way: How Listeria actin subversion has impacted cell biology, infection biology and structural biology. Cell Microbiol. 22, e13190 (2020).

5. R. L. Lamason, M. D. Welch, Actin-based motility and cell-to-cell spread of bacterial pathogens. Curr Opin Microbiol. 35, 48–57 (2017).

6. D. M. Tobin, L. Ramakrishnan, Comparative pathogenesis of Mycobacterium marinum and Mycobacterium tuberculosis. Cell Microbiol. 10, 1027–1039 (2008).

7. A. Aubry, F. Mougari, F. Reibel, E. Cambau, Mycobacterium marinum. Microbiol Spectr. 5 (2017), doi:10.1128/microbiolspec.TNMI7-0038-2016.

8. L. M. Stamm, J. H. Morisaki, L.-Y. Gao, R. L. Jeng, K. L. McDonald, R. Roth, S. Takeshita, J. Heuser, M. D. Welch, E. J. Brown, Mycobacterium marinum escapes from phagosomes and is propelled by actin-based motility. J Exp Med. 198, 1361– 1368 (2003).

9. L. M. Stamm, M. A. Pak, J. H. Morisaki, S. B. Snapper, K. Rottner, S. Lommel, E. J. Brown, Role of the WASP family proteins for Mycobacterium marinum actin tail formation. Proc Natl Acad Sci U S A. 102, 14837–14842 (2005).

10. G. M. Kennedy, J. H. Morisaki, P. A. D. Champion, Conserved mechanisms of Mycobacterium marinum pathogenesis within the environmental amoeba Acanthamoeba castellanii. Appl Environ Microbiol. 78, 2049–2052 (2012).

11. L. S. Ates, A. Dippenaar, R. Ummels, S. R. Piersma, A. D. van der Woude, K. van der Kuij, F. Le Chevalier, D. Mata-Espinosa, J. Barrios-Payán, B. Marquina- Castillo, C. Guapillo, C. R. Jiménez, A. Pain, E. N. G. Houben, R. M. Warren, R. Brosch, R. Hernández-Pando, W. Bitter, Mutations in ppe38 block PE_PGRS secretion and increase virulence of Mycobacterium tuberculosis. Nat Microbiol. 3, 181–188 (2018).

12. A. M. Abdallah, T. Verboom, E. M. Weerdenburg, N. C. Gey van Pittius, P. W. Mahasha, C. Jiménez, M. Parra, N. Cadieux, M. J. Brennan, B. J. Appelmelk, W. Bitter, PPE and PE_PGRS proteins of Mycobacterium marinum are transported via the type VII secretion system ESX-5. Mol Microbiol. 73, 329–340 (2009).

13. M. J. Burggraaf, A. Speer, A. S. Meijers, R. Ummels, A. M. van der Sar, K. V. Korotkov, W. Bitter, C. Kuijl, Type VII Secretion Substrates of Pathogenic Mycobacteria Are Processed by a Surface Protease. mBio. 10 (2019), doi:10.1128/mBio.01951-19.

14. J. A. Olzmann, P. Carvalho, Dynamics and functions of lipid droplets. Nat Rev Mol Cell Biol. 20, 137–155 (2019).

15. M. Bosch, M. Sánchez-Álvarez, A. Fajardo, R. Kapetanovic, B. Steiner, F. Dutra, L. Moreira, J. A. López, R. Campo, M. Marí, F. Morales-Paytuví, O. Tort, A. Gubern, R. M. Templin, J. E. B. Curson, N. Martel, C. Català, F. Lozano, F. Tebar, C. Enrich, J. Vázquez, M. A. Del Pozo, M. J. Sweet, P. T. Bozza, S. P. Gross, R. G. Parton, A. Pol, Mammalian lipid droplets are innate immune hubs integrating cell metabolism and host defense. Science. 370, eaay8085 (2020).

16. R. Gautier, D. Douguet, B. Antonny, G. Drin, HELIQUEST: a web server to screen sequences with specific alpha-helical properties. Bioinformatics. 24, 2101–2102 (2008).

17. M. J. Brennan, G. Delogu, Y. Chen, S. Bardarov, J. Kriakov, M. Alavi, W. R. Jacobs, Evidence that mycobacterial PE_PGRS proteins are cell surface constituents that influence interactions with other cells. Infect Immun. 69, 7326– 7333 (2001).

18. G. Delogu, C. Pusceddu, A. Bua, G. Fadda, M. J. Brennan, S. Zanetti, Rv1818c- encoded PE_PGRS protein of Mycobacterium tuberculosis is surface exposed and influences bacterial cell structure. Mol Microbiol. 52, 725–733 (2004).

19. A. Cascioferro, G. Delogu, M. Colone, M. Sali, A. Stringaro, G. Arancia, G. Fadda, G. Palù, R. Manganelli, PE is a functional domain responsible for protein translocation and localization on mycobacterial cell wall. Mol Microbiol. 66, 1536– 1547 (2007).

20. S. Basu, S. K. Pathak, A. Banerjee, S. Pathak, A. Bhattacharyya, Z. Yang, S. Talarico, M. Kundu, J. Basu, Execution of macrophage apoptosis by PE_PGRS33 of Mycobacterium tuberculosis is mediated by Toll-like receptor 2-dependent release of tumor necrosis factor-alpha. J Biol Chem. 282, 1039–1050 (2007).

21. I. Palucci, S. Camassa, A. Cascioferro, M. Sali, S. Anoosheh, A. Zumbo, M. Minerva, R. Iantomasi, F. De Maio, G. Di Sante, F. Ria, M. Sanguinetti, G. Palù, M. J. Brennan, R. Manganelli, G. Delogu, PE_PGRS33 Contributes to Mycobacterium tuberculosis Entry in Macrophages through Interaction with TLR2. PLoS One. 11, e0150800 (2016).

22. L. S. Ates, R. Ummels, S. Commandeur, R. van de Weerd, R. van der Weerd, M. Sparrius, E. Weerdenburg, M. Alber, R. Kalscheuer, S. R. Piersma, A. M. Abdallah, M. Abd El Ghany, A. M. Abdel-Haleem, A. Pain, C. R. Jiménez, W. Bitter, E. N. G. Houben, Essential Role of the ESX-5 Secretion System in Outer Membrane Permeability of Pathogenic Mycobacteria. PLoS Genet. 11, e1005190 (2015).

23. S. Jäger, P. Cimermancic, N. Gulbahce, J. R. Johnson, K. E. McGovern, S. C. Clarke, M. Shales, G. Mercenne, L. Pache, K. Li, H. Hernandez, G. M. Jang, S. L. Roth, E. Akiva, J. Marlett, M. Stephens, I. D’Orso, J. Fernandes, M. Fahey, C. Mahon, A. J. O’Donoghue, A. Todorovic, J. H. Morris, D. A. Maltby, T. Alber, G. Cagney, F. D. Bushman, J. A. Young, S. K. Chanda, W. I. Sundquist, T. Kortemme, R. D. Hernandez, C. S. Craik, A. Burlingame, A. Sali, A. D. Frankel, N. J. Krogan, Global landscape of HIV-human protein complexes. Nature. 481, 365–370 (2011).

24. M. Symons, J. M. Derry, B. Karlak, S. Jiang, V. Lemahieu, F. Mccormick, U. Francke, A. Abo, Wiskott-Aldrich syndrome protein, a novel effector for the GTPase CDC42Hs, is implicated in actin polymerization. Cell. 84, 723–734 (1996).

25. R. Kolluri, K. F. Tolias, C. L. Carpenter, F. S. Rosen, T. Kirchhausen, Direct interaction of the Wiskott-Aldrich syndrome protein with the GTPase Cdc42. Proc Natl Acad Sci U S A. 93, 5615–5618 (1996).

26. F. De Maio, R. Berisio, R. Manganelli, G. Delogu, PE_PGRS proteins of Mycobacterium tuberculosis: A specialized molecular task force at the forefront of host-pathogen interaction. Virulence. 11, 898–915 (2020).

27. L. S. Ates, New insights into the mycobacterial PE and PPE proteins provide a framework for future research. Mol Microbiol. 113, 4–21 (2020).

28. C. Barisch, T. Soldati, Mycobacterium marinum Degrades Both Triacylglycerols and Phospholipids from Its Dictyostelium Host to Synthesize Its Own Triacylglycerols and Generate Lipid Inclusions. PLoS Pathog. 13, e1006095 (2017).

29. D. M. Tobin, J. C. Vary, J. P. Ray, G. S. Walsh, S. J. Dunstan, N. D. Bang, D. A. Hagge, S. Khadge, M.-C. King, T. R. Hawn, C. B. Moens, L. Ramakrishnan, The lta4h locus modulates susceptibility to mycobacterial infection in zebrafish and humans. Cell. 140, 717–730 (2010).

30. M. Knight, J. Braverman, K. Asfaha, K. Gronert, S. Stanley, Lipid droplet formation in Mycobacterium tuberculosis infected macrophages requires IFN-γ/HIF-1α signaling and supports host defense. PLoS Pathog. 14, e1006874 (2018).

31. S. L. Sampson, Mycobacterial PE/PPE proteins at the host-pathogen interface. Clin Dev Immunol. 2011, 497203 (2011).

32. S. Fishbein, N. van Wyk, R. M. Warren, S. L. Sampson, Phylogeny to function: PE/PPE protein evolution and impact on Mycobacterium tuberculosis pathogenicity. Mol Microbiol. 96, 901–916 (2015).

33. N. K. Saini, A. Baena, T. W. Ng, M. M. Venkataswamy, S. C. Kennedy, S. Kunnath-Velayudhan, L. J. Carreño, J. Xu, J. Chan, M. H. Larsen, W. R. Jacobs, S. A. Porcelli, Suppression of autophagy and antigen presentation by Mycobacterium tuberculosis PE_PGRS47. Nat Microbiol. 1, 16133 (2016).

34. E. J. Strong, T. W. Ng, S. A. Porcelli, S. Lee, Mycobacterium tuberculosis PE_PGRS20 and PE_PGRS47 Proteins Inhibit Autophagy by Interaction with Rab1A. mSphere, e0054921 (2021).

35. V. K. Singh, L. Berry, A. Bernut, S. Singh, S. Carrère-Kremer, A. Viljoen, L. Alibaud, L. Majlessi, R. Brosch, V. Chaturvedi, J. Geurtsen, M. Drancourt, L. Kremer, A unique PE_PGRS protein inhibiting host cell cytosolic defenses and sustaining full virulence of Mycobacterium marinum in multiple hosts. Cell Microbiol. 18, 1489–1507 (2016).

36. Q. Chai, X. Wang, L. Qiang, Y. Zhang, P. Ge, Z. Lu, Y. Zhong, B. Li, J. Wang, L. Zhang, D. Zhou, W. Li, W. Dong, Y. Pang, G. F. Gao, C. H. Liu, A Mycobacterium tuberculosis surface protein recruits ubiquitin to trigger host xenophagy. Nat Commun. 10, 1973 (2019).

37. S. Grover, T. Sharma, Y. Singh, S. Kohli, M. P. A. Singh, T. Semmler, L. H. Wieler, K. Tedin, N. Z. Ehtesham, S. E. Hasnain, The PGRS Domain of Mycobacterium tuberculosis PE_PGRS Protein Rv0297 Is Involved in Endoplasmic Reticulum Stress-Mediated Apoptosis through Toll-Like Receptor 4. mBio. 9, e01017–18 (2018).

38. K. Bansal, S. R. Elluru, Y. Narayana, R. Chaturvedi, S. A. Patil, S. V. Kaveri, J. Bayry, K. N. Balaji, PE_PGRS antigens of Mycobacterium tuberculosis induce maturation and activation of human dendritic cells. J Immunol. 184, 3495–3504 (2010).

39. H. Miki, T. Sasaki, Y. Takai, T. Takenawa, Induction of filopodium formation by a WASP-related actin-depolymerizing protein N-WASP. Nature. 391, 93–96 (1998).

40. R. Rohatgi, L. Ma, H. Miki, M. Lopez, T. Kirchhausen, T. Takenawa, M. W. Kirschner, The interaction between N-WASP and the Arp2/3 complex links Cdc42- dependent signals to actin assembly. Cell. 97, 221–231 (1999).

41. A. S. Kim, L. T. Kakalis, N. Abdul-Manan, G. A. Liu, M. K. Rosen, Autoinhibition and activation mechanisms of the Wiskott-Aldrich syndrome protein. Nature. 404, 151–158 (2000).

42. H.-C. Cheng, B. M. Skehan, K. G. Campellone, J. M. Leong, M. K. Rosen, Structural mechanism of WASP activation by the enterohaemorrhagic E. coli effector EspF(U). Nature. 454, 1009–1013 (2008).

43. N. A. Sallee, G. M. Rivera, J. E. Dueber, D. Vasilescu, R. D. Mullins, B. J. Mayer, W. A. Lim, The pathogen protein EspF(U) hijacks actin polymerization using mimicry and multivalency. Nature. 454, 1005–1008 (2008).

44. R. P. M. Mauricio, C. M. Jeffries, D. I. Svergun, J. E. Deane, The Shigella Virulence Factor IcsA Relieves N-WASP Autoinhibition by Displacing the Verprolin Homology/Cofilin/Acidic (VCA) Domain. J Biol Chem. 292, 134–145 (2017).

45. C. Egile, T. P. Loisel, V. Laurent, R. Li, D. Pantaloni, P. J. Sansonetti, M. F. Carlier, Activation of the CDC42 effector N-WASP by the Shigella flexneri IcsA protein promotes actin nucleation by Arp2/3 complex and bacterial actin-based motility. J Cell Biol. 146, 1319–1332 (1999).

46. R. Faris, A. McCullough, S. E. Andersen, T. O. Moninger, M. M. Weber, The Chlamydia trachomatis secreted effector TmeA hijacks the N-WASP-ARP2/3 actin remodeling axis to facilitate cellular invasion. PLoS Pathog. 16, e1008878 (2020).

47. H. N. Higgs, T. D. Pollard, Activation by Cdc42 and PIP(2) of Wiskott-Aldrich syndrome protein (WASp) stimulates actin nucleation by Arp2/3 complex. J Cell Biol. 150, 1311–1320 (2000).

48. V. Papayannopoulos, C. Co, K. E. Prehoda, S. Snapper, J. Taunton, W. A. Lim, A polybasic motif allows N-WASP to act as a sensor of PIP(2) density. Mol Cell. 17, 181–191 (2005).

49. D. Szklarczyk, A. L. Gable, K. C. Nastou, D. Lyon, R. Kirsch, S. Pyysalo, N. T. Doncheva, M. Legeay, T. Fang, P. Bork, L. J. Jensen, C. von Mering, The STRING database in 2021: customizable protein-protein networks, and functional characterization of user-uploaded gene/measurement sets. Nucleic Acids Res. 49, D605–D612 (2021).

50. C. Prévost, M. E. Sharp, N. Kory, Q. Lin, G. A. Voth, R. V. Farese, T. C. Walther, Mechanism and Determinants of Amphipathic Helix-Containing Protein Targeting to Lipid Droplets. Dev Cell. 44, 73–86.e4 (2018).

51. L. Ramakrishnan, S. Falkow, Mycobacterium marinum persists in cultured mammalian cells in a temperature-restricted fashion. Infect Immun. 62, 3222–3229 (1994).

52. R. Goude, T. Parish, Electroporation of mycobacteria. Methods Mol Biol. 465, 203– 215 (2009).

53. T. P. Burke, P. Engström, R. A. Chavez, J. A. Fonbuena, R. E. Vance, M. D. Welch, Inflammasome-mediated antagonism of type I interferon enhances Rickettsia pathogenesis. Nat Microbiol. 5, 688–696 (2020).

54. H. W. Johnson, M. J. Schell, Neuronal IP3 3-Kinase is an F-actin–bundling Protein: Role in Dendritic Targeting and Regulation of Spine Morphology. Mol Biol Cell. 20, 5166–5180 (2009).

55. R. L. Lamason, E. Bastounis, N. M. Kafai, R. Serrano, J. C. Del Álamo, J. A. Theriot, M. D. Welch, Rickettsia Sca4 Reduces Vinculin-Mediated Intercellular Tension to Promote Spread. Cell. 167, 670–683.e10 (2016).

56. C. A. Schneider, W. S. Rasband, K. W. Eliceiri, NIH Image to ImageJ: 25 years of Image Analysis. Nat Methods. 9, 671–675 (2012).

57. C. M. Sassetti, D. H. Boyd, E. J. Rubin, Genes required for mycobacterial growth defined by high density mutagenesis. Mol Microbiol. 48, 77–84 (2003).

58. M. S. Siegrist, E. J. Rubin, Phage transposon mutagenesis. Methods Mol Biol. 465, 311–323 (2009).

59. D. Barkan, C. L. Stallings, M. S. Glickman, An improved counterselectable marker system for mycobacterial recombination using galK and 2-deoxy-galactose. Gene. 470, 31–36 (2011).

60. C. K. Stover, V. F. de la Cruz, T. R. Fuerst, J. E. Burlein, L. A. Benson, L. T. Bennett, G. P. Bansal, J. F. Young, M. H. Lee, G. F. Hatfull, New use of BCG for recombinant vaccines. Nature. 351, 456–460 (1991).

61. A. Parikh, D. Kumar, Y. Chawla, K. Kurthkoti, S. Khan, U. Varshney, V. K. Nandicoori, Development of a new generation of vectors for gene expression, gene replacement, and protein-protein interaction studies in mycobacteria. Appl Environ Microbiol. 79, 1718–1729 (2013).

62. B. H. Penn, Z. Netter, J. R. Johnson, J. Von Dollen, G. M. Jang, T. Johnson, Y. M. Ohol, C. Maher, S. L. Bell, K. Geiger, G. Golovkine, X. Du, A. Choi, T. Parry, B. C. Mohapatra, M. D. Storck, H. Band, C. Chen, S. Jäger, M. Shales, D. A. Portnoy, R. Hernandez, L. Coscoy, J. S. Cox, N. J. Krogan, An Mtb-Human Protein-Protein Interaction Map Identifies a Switch between Host Antiviral and Antibacterial Responses. Mol Cell. 71, 637–648.e5 (2018).

63. K. R. Clauser, P. Baker, A. L. Burlingame, Role of accurate mass measurement (+/- 10 ppm) in protein identification strategies employing MS or MS/MS and database searching. Anal Chem. 71, 2871–2882 (1999).

64. L. K. Doolittle, M. K. Rosen, S. B. Padrick, Measurement and analysis of in vitro actin polymerization. Methods Mol Biol. 1046, 273–293 (2013).

